# MANY PATHS TO DESTRUCTION: FAMILY-SPECIFIC TURNOVER AND STRESS RESPONSES FOR TRNA INTRONS

**DOI:** 10.1101/2025.11.19.689297

**Authors:** Sara Metcalf, Regina T. Nostramo, Alicia Bao, Katherine Clark, Paolo L. Sinopoli, Anita K. Hopper

## Abstract

In organisms ranging from Archaea to humans, a subset of genes encoding tRNAs contain introns. Upon splicing, the tRNA exons are joined and the released free introns are rapidly degraded. Although tRNAs introns were previously considered to be “junk” sequences, we recently reported that free tRNA introns (fitRNAs) of *S. cerevisiae* serve as negative regulators of the cellular levels of mRNAs that bear long stretches of open reading frame sequence complementarity to tRNA introns. We also reported that 2 of the 10 families of tRNA introns, accumulate to elevated levels when cells suffer oxidative stress. The results led to the current investigations of the regulation of tRNA intron cellular levels. We document that tRNA intron turnover occurs by combinations of 5’ RNA kinases, 5’ to 3’ and 3’ to 5’ exonucleases as well as by at least three endonucleases and, generally, the levels of each tRNA intron family are regulated by a unique combination of nucleases/kinases. Similarly, one family of excised intron can form circles whereas the other free tRNA intron families do not. Further, levels of individual tRNA introns differ in response to environmental conditions including type of media, stage in growth curves, and exposure to elevated temperature. Together, these findings highlight the many cellular pathways utilized to regulate tRNA intron levels and the specificity of these pathways for different tRNA families and varying cellular conditions. The results underscore the likely important roles of the newly discovered individual fitRNAs in regulation of cell biology and responses to environmental conditions.

## INTRODUCTION

The genomes of nearly all archaea and eukaryotic cells encode a subset of tRNA genes which possess introns (1, 2). Although tRNA genes are redundant in genomes, generally if one member of a given tRNA isodecoder family contains an intron then all of the reiterated copies of this family will also encode an intron and for yeast the reiterated sequences are identical or near identical (1). The short (∼ ≥10 and ≤100 nt) intron sequences are located in precursor tRNA (pre-tRNA) transcripts one nucleotide 3’ to the anticodon, interrupting the tRNA anticodon stem-loop [Reviews (3, 4)]. Thus, pre-tRNAs containing introns do not participate in protein synthesis and consequently, removal of intron sequences from intron-containing pre-tRNAs via tRNA splicing is essential for decoding the genome into the proteome.

Pre-tRNA splicing in eukaryotes is catalyzed by the essential heterotetrameric tRNA splicing endonuclease complex (SEN in yeast; TSEN in vertebrates) (4, 5). Splicing generates a 5’ exon with a 2’, 3’ cyclic phosphate, a 3’ exon bearing a 5’ hydroxyl group, and a free tRNA intron (4–6). The endolytic splicing mechanism is highly conserved from archaea to vertebrates (4, 7, 8). Subsequent to intron removal the 5’ and 3’ exons are spliced together by tRNA ligases. In archaea, *Drosophila*, and vertebrates the RtcB ligase reaction resolves the 2’, 3’ cyclic phosphate and directly joins the resulting 3’ phosphate of the 5’ exon with the 5’ hydroxyl of the 3’ exon (6, 9). In contrast, in yeast, protozoa, and plants, ligation proceeds via a more complicated reaction catalyzed by a multidomain protein, Rlg1/Trl1 (hereafter referred to as Trl1), and Tpt1 (4, 9–11). Except for recently discovered ligases from the fungal order Mucorales, Trl1 enzymes possess three activities in single polypeptides (4, 9, 12, 13): (1) opening of the 2’, 3’ cyclic phosphate of the 5’ exon by the Trl1 C-terminal cyclic phosphodiesterase (CPD) domain, generating a linear 5’ exon bearing a 3’ hydroxyl and a 2’ phosphate; (2) phosphorylation of the 5’ hydroxyl of the 3’ exon catalyzed by the Trl1 RNA kinase domain; (3) ligation of the 5’ and 3’ exons catalyzed by the Trl1 amino-terminal ligase domain. The ligation step generates a linear spliced RNA bearing two phosphates, one in the polynucleotide backbone and the other in the 2’ position at the splice junction. In the final step of the tRNA splicing reaction the 2’ phosphate at the splice junction is removed in a phosphotransferase reaction by Tpt1 (4, 11).

Completion of the pre-tRNA splicing reaction generates mature tRNAs and freed introns. Interestingly, the free introns from *Drosophila* pre-tRNAs form covalent circles (6, 14), whereas introns from budding yeast are linear molecules (4, 5, 15). Not only is possession of introns in subsets of tRNA genes conserved, but, in addition, for *Drosophila* and budding yeast there is phylogenetic conservation of intron sequences among tRNA isodecoder family members (16, 17). Despite this conservation, until recently, freed tRNA introns were not known to serve any biological function. However, we recently showed that the free tRNA^Ile^_UAU_ and tRNA^Trp^_CCA_ introns from *S. cerevisiae* serve as negative regulators of the levels of mRNAs bearing extended complementary sequences in the open reading frames (ORFs) to the free introns (17), thereby regulating basal levels of the mRNAs. We also demonstrated that yeast cells exposed to H_2_O_2_ stress accumulate elevated levels of tRNA^Trp^_CCA_ and tRNA^Leu^_CAA_ introns, but do not accumulate elevated levels of the remaining 8 families of tRNA introns (17). Thus, in contradiction to the notion that free tRNA introns are junk to be rapidly disposed of, we discovered that free tRNA introns (fitRNAs) serve to regulate both basal and stress-induced gene expression. Therefore, it is important to understand the mechanisms by which free tRNA intron cellular levels are regulated. Here, we report that for the studied yeast free tRNA intron families, intron cellular levels are determined by a myriad of tRNA family-specific nucleases/kinases as well as by specific growth conditions/stresses. The results underscore likely important roles of individual tRNA intron sequences in regulation of cell biology and responses to environmental conditions.

## RESULTS

### Roles of 5’ to 3’ exonuclease Xrn1 and Trl1 ligase in family-specific tRNA intron turnover

Even though eukaryotic cells generate enormous quantities of free tRNA introns per cell cycle (4), free tRNA introns are rarely detected in RNA samples isolated from cells grown under normal physiological conditions. This is because tRNA introns are rapidly and efficiently destroyed. We discovered one pathway that degrades budding yeast linear tRNA^Ile^_UAU_ intron (15). This linear 60 nt RNA is turned over in a two-step process. In step 1, the RNA kinase activity of tRNA ligase Trl1 phosphorylates the 5’ hydroxyl of the intron. In step 2, the resulting 5’ monophosphate renders the intron as a substrate for the cytoplasmic 5’ to 3’ exonuclease, Xrn1, which efficiently destroys the tRNA^Ile^_UAU_ intron (15, 18).

Ten families of *S. cerevisiae* tRNAs are encoded by genes containing introns. We investigated whether the same 2-step mechanism elucidated for tRNA^Ile^_UAU_ intron turnover is employed for the other families. Employing a nonradioactive northern blotting procedure (10, 19) and tRNA intron-specific probes (Supplementary Table 1), we assessed the levels of free tRNA introns in RNAs isolated from wild-type (WT) cells, cells harboring an *XRN1* deletion (*xrn1Δ*), and cells with temperature sensitive mutations of the essential *TRL1* gene [*trl1-4* (10) or *trl1-ts* (20)]. The strains were grown in rich (YEPD) media at 23°C, then shifted to 37°C for 2 hr prior to isolation of small RNAs and subsequent northern blot analysis (19).

We anticipated that cellular levels of each of the families of tRNA introns would be dependent upon the same 2-step turnover mechanism as had been elucidated for the tRNA^Ile^_UAU_ intron. Contrary to expectations, only the tRNA^Ile^_UAU_ and tRNA^Leu^_CAA_ introns possess elevated levels in both *xrn1Δ* and *trl1-ts* mutant cells (Fig. 1). The tRNA^Trp^_CCA_ and tRNA^Pro^_UGG_ introns accumulated at levels higher than WT cells in the *xrn1Δ* mutant, but not in *trl1* mutant cells (Fig. 1). Since Xrn1 primarily degrades RNA substrates bearing 5’ monophosphates (18), this Xrn1-dependent, Trl1-independent turnover mechanism employs an unknown RNA kinase. Further, the cellular levels of introns from tRNA^Lys^_UUU_, tRNA^Phe^_GAA_, and tRNA^Ser^_GCU_ did not accumulate in *xrn111* cells above the levels in WT cells but did accumulate in RNAs isolated from *trl1* mutant cells. Thus, turnover of tRNA^Lys^_UUU_, tRNA^Phe^_GAA_, and tRNA^Ser^_GCU_ introns is Xrn1-independent and Trl1-dependent (Fig. 1); presumably, 5’ phosphorylation of these three families renders their turnover by a 5’ to 3’ exonuclease other than Xrn1. Surprisingly, neither tRNA^Ser^_CGA_ nor tRNA^Leu^_UAG_ intron levels were altered by mutation of *XRN1* or *TRL1*; so, turnover of tRNA^Ser^_CGA_ and tRNA^Leu^_UAG_ introns is both Xrn1- and Trl1-independent (Fig. 1). Finally, even though our probe for the 14 nucleotide tRNA^Tyr^_GUA_ intron sequence hybridized efficiently to the initial and end-processed pre-tRNAs, we were unable to detect the free tRNA^Tyr^_GUA_ intron by northern analysis (Fig. 1); so, dependence upon Trl1 and Xrn1 for tRNA^Tyr^_GUA_ intron turnover remains unknown.

**Figure 1.**
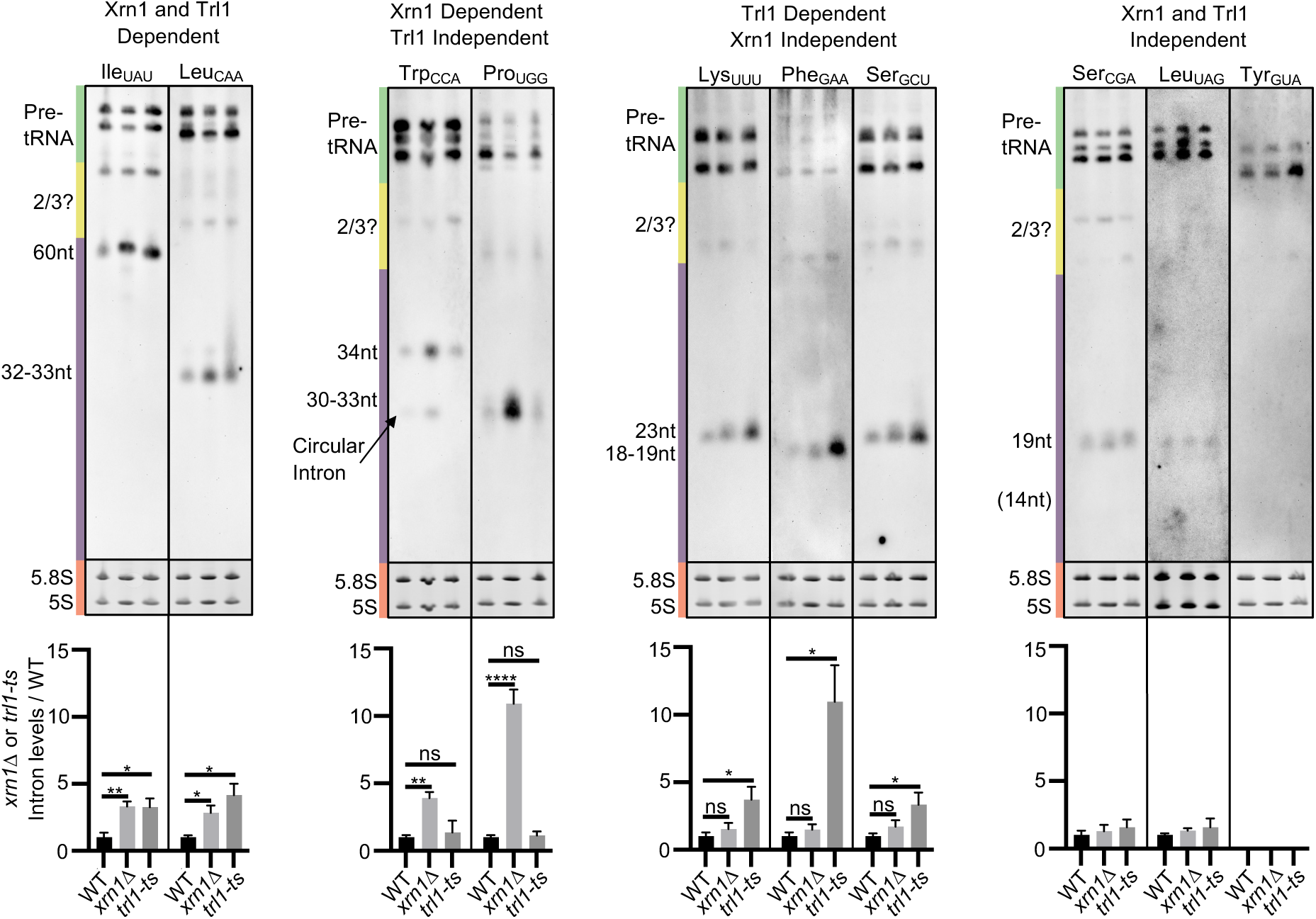
Different roles of Xrn1 and Trl1 in turnover of individual tRNA intron families: Wild-type (WT), *xrn111*, and *trl1-ts* cells were grown in YEPD media to early log phase and then shifted to 37°C, the non-permissive temperature for *trl1-ts*) for 2 hr prior to harvesting cells by centrifugation. “Small” RNAs were extracted, resolved on polyacrylamide gels and analyzed by northern blotting employing digoxigenin-labeled oligonucleotides complementary to the indicated tRNA intron (Supplementary Table 1). For these full-length blots, the levels of initial tRNA transcripts and end-processed intron-containing pre-tRNA are indicated by a green bar; putative 2/3 tRNA containing the tRNA intron and an exon are indicated by a yellow bar; and the free intron is indicated by a purple bar. EtBr staining of 5.8S and 5S rRNAs (orange bar) serve as loading controls. The arrow indicates the tRNA^Trp^_CCA_ circular intron. Quantitation of intron levels is displayed below, expressed as intron levels relative to 5S rRNA levels in mutant cells vs. WT cells (set as 1). Data, are expressed as mean ± SEM. *n* ≥ 3, with a single representative northern blot shown. \**p* < 0.05; \*\**p* < 0.01; \*\*\**p* < 0.001; \*\*\*\**p* < 0.0001; ns: p > 0.05.

Our studies of the dependence of free tRNA intron levels upon functional Xrn1 and Trl1 document that there are at least 4 different family-specific turnover mechanisms for free tRNA introns. However, further inspection of the tRNA^Trp^_CCA_ intron uncovered additional complexities. The probe specific to the tRNA^Trp^_CCA_ intron detects two differently migrating RNAs isolated from *xrn111* cells; the predominant slower RNA migrates as expected for a linear 34 polynucleotide, whereas the less prominent RNA migrates somewhat faster than expected (Fig. 1). In addition to aberrant migration on polyacrylamide gels our studies documented that the faster migrating tRNA^Trp^_CCA_ intron is a covalent circle. First, to distinguish whether the aberrantly migrating RNA is linear or circular we treated RNA isolated from WT and *xrn111* cells with TEX, a 5’ to 3’ RNase, which efficiently degrades linear RNAs bearing a 5’ monophosphate (21). Migration of 5S rRNA which is resistant to TEX digestion, likely due to its secondary structure and 5’ triphosphate, served as the loading control and 5.8S rRNA, previously shown to be efficiently degraded by TEX (15), served as the positive control. As anticipated for a linear RNA, the slower migrating tRNA^Trp^_CCA_ intron, which is prominent in untreated RNAs extracted from *xrn111* cells, was nearly undetectable upon TEX treatment; in contrast, the faster migrating tRNA^Trp^_CCA_ intron was resistant to TEX treatment (Fig. 2A) as would be expected for a circular RNA lacking a 5’ terminus. Secondly, the levels of the faster migrating tRNA^Trp^_CCA_ intron are dependent upon Trl1 activity as it is barely detected in RNAs isolated from the *trl1-ts* mutant strain incubated at 37°C for 2 hr (Fig. 2B). Although excised tRNA introns from *Drosophila* are circularized via a direct ligation mechanism by the tRNA ligase, RtcB (16) and yeast tRNA introns can be circularized when provided with RtcB *in vivo* (14), prior studies have reported all yeast excised tRNA introns to be linear (5, 15). Why among the 8 studied yeast tRNA introns only the tRNA^Trp^_CCA_ intron exists *in vivo* as both linear and circular molecules is unknown, but turnover of this circular form of a tRNA intron would require an endonuclease, rather than an exonuclease, implicating a 5^th^ mechanism for family-specific tRNA intron turnover.

**Figure 2.**
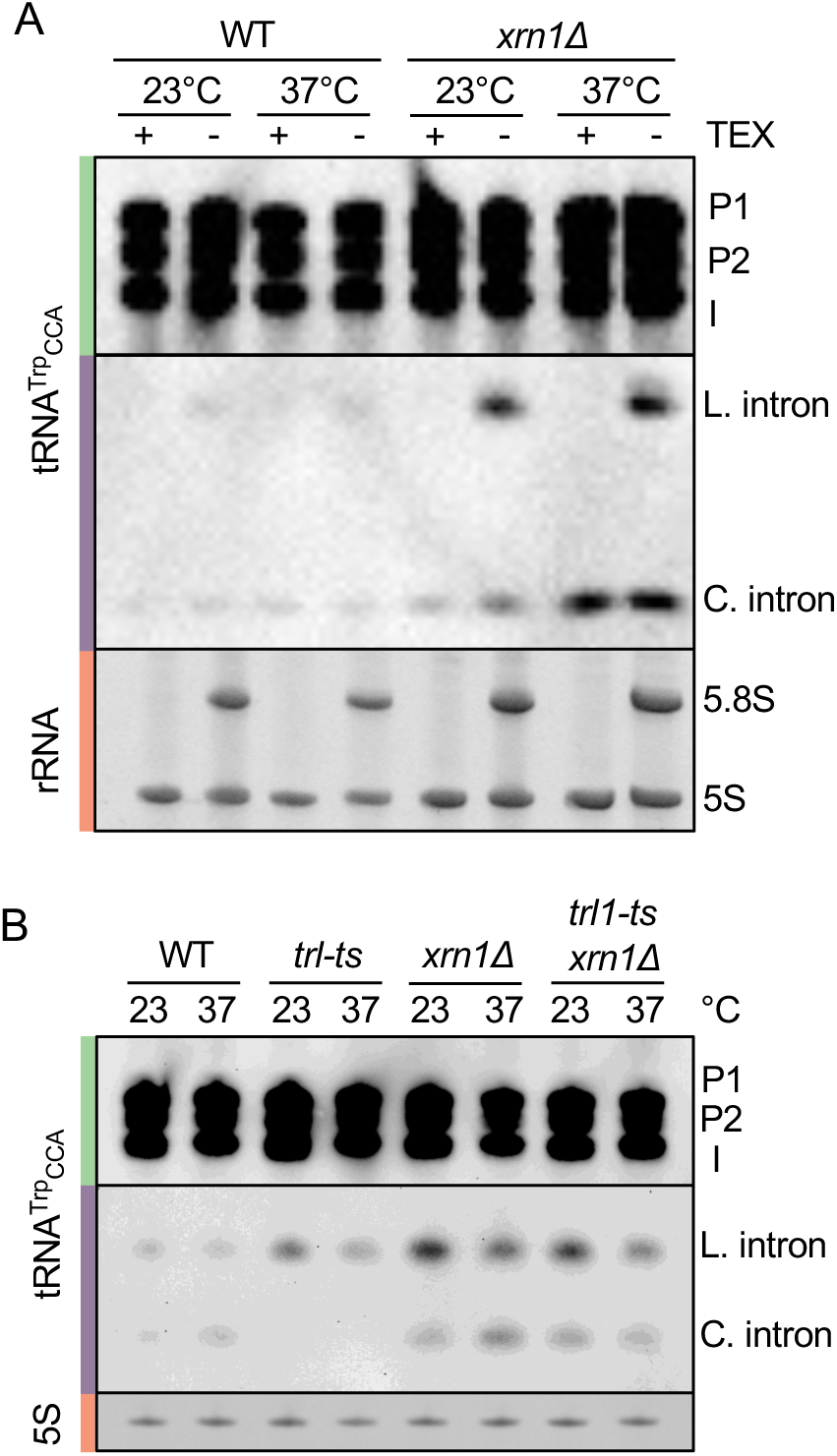
The tRNA^Trp^_CCA_ intron exists in both linear and circular forms: **A.** WT *or xrn111* cells were grown in YEPD to early log phase at either 23^0^C or 37^0^C for 2 hr. Small RNAs were isolated, treated with the 5’→3’ exonuclease, TEX, and then subjected to northern blot analyses. The green bar depicts the pre-tRNAs: P1 is the initial transcript; P2 is the 5’ leader-containing, 3’ end-processed intron-containing tRNA (60); I is the 5’ and 3’ end-processed intron-containing tRNA. The purple bar delineates the tRNA intron; the slower and faster migrating forms are indicated as L. and C., respectively. The orange bar depicts 5.8S and 5S rRNAs as detected by EtBr staining. **B**. Circularization of the tRNA^Trp^_CCA_ intron requires Trl1 activity. RNAs isolated from WT, *trl1-ts, xrn111, or trl1-ts xrn111* cells grown in YEPD to early log phase at either 23^0^C or after shift to 37^0^C for 2 hr. Northern blots are as described for A. 5S rRNA serves as the loading control.

Further analyses of the tRNA^Leu^_CAA_ intron also uncovered complexities as its migration differed depending on the *trl1* allele (*trl1-4* vs. *trl1-ts*). The *trl1-4* allele has a single nucleotide change, T180I, located in the ligase domain [(10); (Fig. 3A)]. Although the T180I change is located in the ligase domain, our previous studies showed that the mutant enzyme is also defective in kinase activity (15). A second *trl1* temperature sensitive allele (*trl1-ts*) was generated by the Hieter lab (20). We sequenced this mutant gene and surprisingly found that it possesses 12 nucleotide changes; 7 mutations are located in the ligase domain, 3 map to the kinase domain, and 2 map to the CPD domain (Fig. 3A; Supplementary Fig. 1). With exception of the tRNA^Leu^_CAA_ intron, RNAs isolated from *trl1-4* and *trl1-ts* strains possess the same tRNA intron accumulation patterns (Fig. 3B). Although the tRNA^Leu^_CAA_ intron migrates as a prominent single and less prominent slower banding pattern from RNAs isolated from the *trl1-4* strain, the tRNA^Leu^_CAA_ intron from RNAs isolated from the *trl1-ts* “smears” upward with some “laddering” (Figs. 1 and 3B). The aberrantly migrating tRNA^Leu^_CAA_ was not altered upon in vitro digestion with TEX (Supplementary Fig. 2), supporting that it lacks an accessible 5’ phosphate. The results indicate that different *trl1* alleles cause unanticipated differing consequences upon the turnover/structure of the tRNA^Leu^_CAA_ intron.

**Figure 3.**
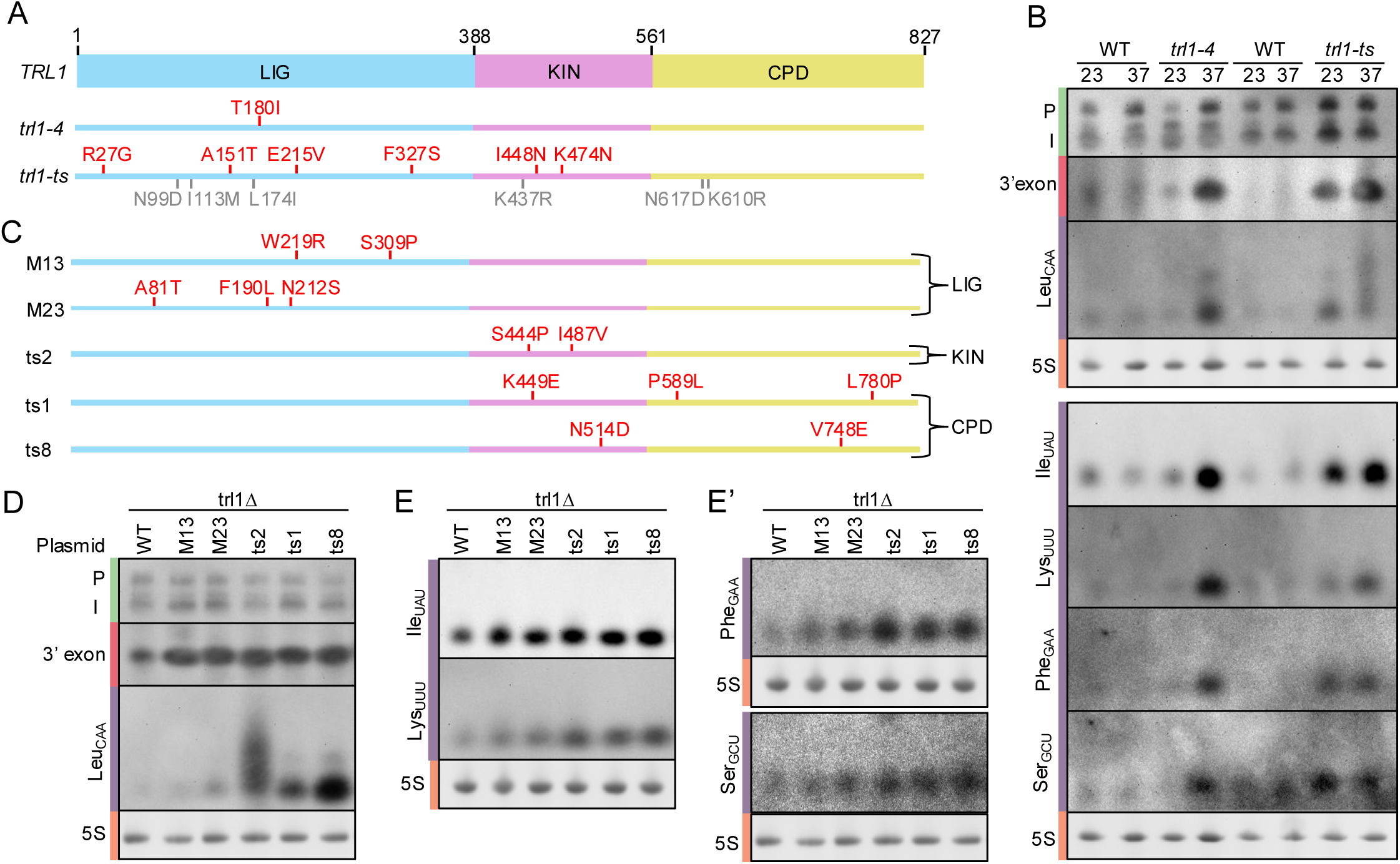
Differing roles for the three Trl1 enzyme activities in tRNA intron turnover. **A**. Diagram depicting Trl1 enzyme activity domains: Blue -ligase (LIG) domain; pink - kinase (KIN) domain; and yellow - cyclic phosphodiesterase (CPD) domain. The locations and identity of mutations in *trl1-4* (10) vs *trl1-ts* [(20); see methods] alleles are indicated in red font for mutations likely to affect protein structure/function and grey font for mutations unlikely to do so. **B.** Only the tRNA^Leu^_CAA_ intron migrates differently in RNAs extracted from *trl1-4* vs. *trl1-ts* mutant cells. Northern blot analyses showing tRNA intron levels in RNAs isolated from WT (SS330) and *trl1-4* vs WT (BY4741) and *trl1-ts* strains grown in YEPD at 23°C or harvested after a 2 hr shift to 37^0^C. Blots were probed with oligonucleotides complementary to the indicated tRNA intron or the tRNA^Leu^_CAA_ 5’ exon as indicated. **C.** Diagram depicting the locations of *TRL1* temperature sensitive mutations of plasmid-encoded *trl1* alleles. The M13 and M23 plasmids have defects in ligase activity; the ts2 plasmid has defects in kinase activity, and the ts1 and ts8 plasmids have defects in CPD activity [(22); also see Ramirez 2009 cited in the text]. **D, E,** and **E’.** Analysis of the roles of individual Trl1 domains for the five Trl1-dependent tRNA introns. *trl1Δ* cells containing either a WT or temperature-sensitive mutant Trl1 plasmid, were grown at 23°C, then shifted to 37^0^C for 2 hrs. Small RNAs were isolated and northern blot analysis was performed as in (C). For (C) and (D), pre-tRNAs are indicated by the green bars, the tRNA^Leu^_CAA_ 3’ exon is indicated by the red bar, the free tRNA introns are indicated by the purple bars and 5S rRNA levels by the orange bars. Blots were used for multiple rounds by probing, gently stripping and re-probing the membranes.

To determine which Trl1 domain is responsible for the aberrant migration on polyacrylamide gels, we employed plasmids encoding ts mutations of individual Trl1 domains [Ramirez, A. (2009). *Analysis of the RNA phosphotransferase and tRNA splicing activities of human Clp1* (Order No. 3383875). Available from ProQuest Dissertations & Theses Global. (305132011). https://proxy.lib.ohio-state.edu/login?url=https://www.proquest.com/dissertations-theses/analysis-rna-phosphotransferase-trna-splicing/docview/305132011/se-2)] (22); Supplementary Table 2). The parent strain possesses a lethal *trl1Δ* which was complemented by a plasmid-encoded wild-type *TRL1* gene. Plasmids encoding temperature-sensitive mutations of the ligase, kinase, or cyclic phosphodiesterase domains were substituted for the plasmid encoding the *TRL1* wild-type gene by plasmid shuffling (Fig. 3C). Inactivation of ligase activity for either of two alleles – M13 (W219R, S309P) and M23 (A81T, F190L, N212S) - did not alter the levels nor migration pattern of the tRNA^Leu^_CAA_ intron (Fig. 3D); thus, Trl1 ligase activity apparently is not involved in tRNA^Leu^_CAA_ intron turnover. Surprisingly, inactivation of the kinase activity by the ts2 allele (S444P, I487V) resulted in higher levels of the tRNA^Leu^_CAA_ intron which migrated as a smear with laddering (Fig. 3D), quite similar to the pattern obtained for RNA isolated from the *trl1-ts* allele (Figs. 1, 3B). It remains unknown why the *trl1-4* and *trl1-ts* alleles, both of which affect Trl1 kinase activity, differently affect the migration of tRNA^Leu^_CAA_ on polyacrylamide urea gels. Unexpectedly, inactivation of the cyclic phosphodiesterase activity by the ts1 allele (K449E, P589L, L780P) or the ts8 allele (N514D, V748E) resulted in significant accumulation of the faster migrating tRNA^Leu^_CAA_ intron Fig. 3C); thus, turnover of the tRNA^Leu^_CAA_ intron requires resolution of the 2’,3’ cyclic phosphate bond.

We determined the roles of the individual catalytic domains for the remaining Trl1-dependent tRNA introns and learned that each intron (tRNA^Ile^_UAU_, tRNA^Lys^_UUU_, tRNA^Phe^_GAA_, and tRNA^Ser^_GCU_) accumulated in cells with defects in both the Trl1 kinase and the Trl1 CPD domains and none had aberrant migration, as was observed for the tRNA^Leu^_CAA_ intron (Fig. 3E and E’). An interpretation of the increased levels of tRNA introns from families which are dependent upon Trl1 when CPD activity is ablated is that these tRNA introns may be degraded by 3’ to 5’ exonucleases that are dependent upon decyclization of the 2’,3’ cyclic phosphodiester bond (see below). In summary, the results document that there are multiple, and likely, redundant mechanisms, for turnover of the individual budding yeast tRNA intron families.

### Myriad of proteins and redundant pathways for tRNA intron turnover

The above studies of the roles for Xrn1 and Trl1 in tRNA intron turnover led to the appreciation that unknown nucleases and/or kinases participate in tRNA intron turnover. Our strategy to identify the unknown nucleases and/or kinases was to screen mutants bearing lesions in genes encoding annotated nucleases and kinases for the consequences upon tRNA intron levels. We assessed the levels of 8 of the 10 family-specific tRNA introns in RNAs isolated from numerous yeast mutants harboring deletions or conditional ts mutations. For most studies we eliminated northern analyses of tRNA^Tyr^_GUA_ and tRNA^Ser^_CGA_ introns as the signals of these introns were often not visible, very faint, and/or noisy.

For unessential genes we employed the commercially available *MATa* and *MATα* deletion collections (23) and for essential genes we screened yeast harboring temperature sensitive (ts) alleles from the Boone (24) and Heiter (20) collections. The genes screened (Supplementary Table 3) included mutations of: (i) RNA kinase Grc3 (25) and putative RNA kinase Clp1 whose *Drosophila* orthologue negatively regulates circularization of tRNA introns (26); (ii) 5’ to 3’ nucleases Rat1 (18) and its regulator, Rai1 (27), and Dxo1 (28); (iii) 3’ to 5’ exonucleases Rex2, Rex3, and Rex4, involved in 3’ end processing of snRNAs, rRNAs, and ncRNAs (28); Pop2, the mRNA deadenylation exonuclease of the Ccr4-Not complex (29), and the exosome catalytic enzymes Rrp6 and Dis3 [Review:(30)] as well as the Trf4 and Trf5, the nucleotidyl transferase components of the TRAMP complex [Review (31)], and noncatalytic exosome structural and auxiliary proteins: Mtr3, Ski2, Ski3, Ski7, Ski8, Rrp4, and Rrp45 (30); and (iv) numerous endonucleases/factors including Ire1 (32), Rny1 (33), Las1 (34), Yth1 (35), Ysh1 (36), Nob1(37), Dbr1 (38), and Pop3, Pop4, and Pop7 subunits of the RNase P and MRP complexes [Reviews (39, 40)] and nonspecific endonuclease, Nuc1 (41). Those mutants which did not detectably affect the levels of <u>any</u> of the 8 tRNA introns investigated were eliminated from further consideration (Supplementary Table 3, black font). In contrast, those candidates found to affect levels of <u>at least one</u> of the 8 tested tRNA introns were further verified by complementation (where possible) and/or by studies of independently derived mutants of the candidate genes, and/or by differences in accumulation in permissive vs. nonpermissive temperatures (Supplementary Table 3, red font; Supplementary Fig. 3). The results of the candidate screen uncovered a myriad of proteins and pathways functioning, directly or indirectly, in tRNA intron turnover as detailed below and summarized in Fig. 4 and Supplementary Table 4.

**Figure 4.**
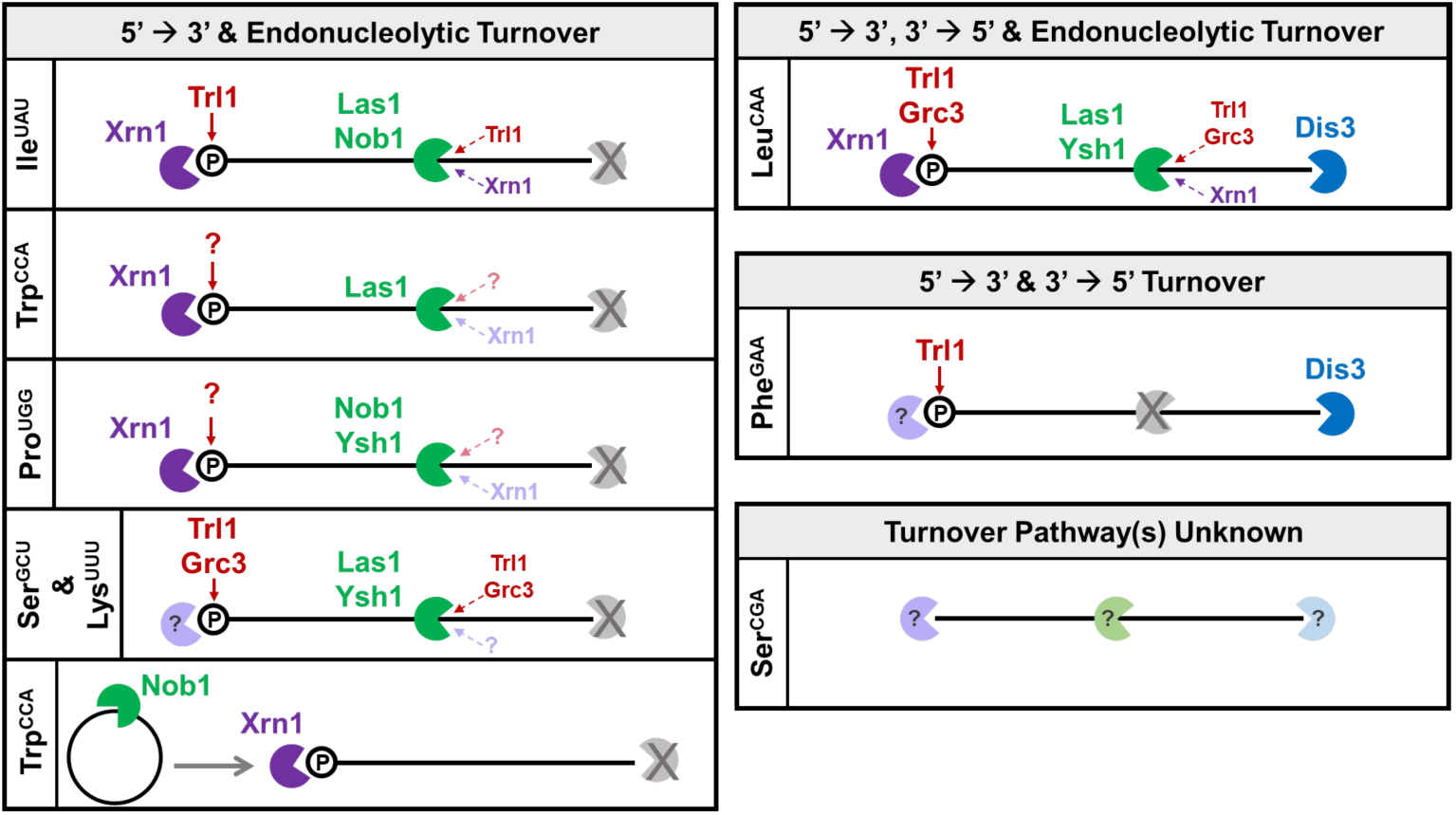
Cartoon depicting the roles of kinases and nucleases in tRNA intron turnover. The introns from each tRNA family are categorized based on whether 5’ to 3’, 3’ to 5’ and/or endonuclease digestion pathways are utilized. 5’ RNA kinases are depicted in red; 5’ to 3’ exonucleases are depicted in purple; 3’ to 5’ exonucleases are depicted in blue; and endonucleases are depicted in green. Unidentified turnover pathways are depicted in grey. For clarity, endonuclease digestion is depicted at a single site in the middle of the intron, with the same site utilized for all identified endonucleases. However, the site(s) of endonucleolytic cleavage is unknown. Question marks are used to depict likely, but currently unidentified turnover components. For example, introns whose turnover is dependent on a 5’ kinase render the intron a substrate for 5’ to 3’ exonucleolytic degradation. Introns whose turnover is dependent on a 5’ to 3’ exonuclease likely require 5’ phosphorylation by an RNA kinase. Dotted red and purple arrows indicate likely (darker) or possible (lighter) 5’ phosphorylation and subsequent 5’ to 3’ degradation, respectively, of the 3’ intron fragment generated by endonucleolytic digestion. It should be noted that cleavage by the endonuclease Nob1 yields a phosphorylated 3’ fragment that could be degraded by Xrn1.

#### 5’ to 3’ turnover of tRNA introns

Neither Rat1 nor Dxo1 mutations affected levels of any of the 8 tested tRNA introns. Since Xrn1 and Rat1 are paralogue RNases which function in different subcellular locations [(Xrn1 is cytoplasmic whereas Rat1 is nuclear (42)], the results could indicate that 5’ to 3’ tRNA intron turnover of the introns dependent upon Xrn1 (tRNA^Ile^_UAU_, tRNA^Leu^_CAA_, tRNA^Trp^_CCA_, and tRNA^Pro^_UGG_) is solely or largely restricted to the cytoplasm as would be expected given that splicing in yeast occurs in the cytoplasm (43, 44). A caveat of this interpretation is that *rat1-1* cells were incubated at 37°C for 2 hr prior to RNA isolation and this could have been insufficient to completely inactivate Rat1.

Xrn1 degrades RNA substrates that bear a 5’ monophosphate (18). As detailed above Trl1 possess 5’ nucleotide kinase activity but Xrn1 turnover of tRNA^Trp^_CCA_ and tRNA^Pro^_UGG_ introns is Trl1 independent. The only other annotated yeast gene encoding 5’ RNA kinase activity besides *TRL1* is *GRC3.* However, neither the tRNA^Trp^_CCA_ nor the tRNA^Pro^_UGG_ introns accumulate in the *grc3-ts* mutant at the nonpermissive temperature (Supplementary Fig. 3A). Thus, if the 5’ to 3’ degradation of the tRNA^Trp^_CCA_ and tRNA^Pro^_UGG_ introns occurs starting from the excised 5’ end or a newly generated 5’ end bearing a 5’ hydroxyl following endonucleolytic cleavage (see endonuclease turnover, below), then 5’ kinase activity required for Xrn1 exonuclease activity remains unidentified. Surprisingly, there is accumulation of other tRNA introns which are Trl1-dependent (tRNA^Leu^_CAA_, tRNA^Lys^_UUU_, and tRNA^Ser^_GCU_) in the *grc3-ts* mutant and the accumulation is complemented upon introduction of a low copy vector encoding WT *GRC3* (Supplementary Fig. 3A), indicating Grc3 RNA kinase redundancy with Trl1 for turnover of these tRNA introns (Fig. 4).

#### 3’ to 5’ tRNA intron turnover of linear tRNA introns

Turnover of the 5 tRNA introns that are dependent upon Trl1 CPD activity [(tRNA^Ile^_UAU_, tRNA^Leu^_CAA_, tRNA^Lys^_UUU_, tRNA^Phe^_GAA_, and tRNA^Ser^_GCU_); (Fig. 3)] might be substrates of 3’ to 5’ RNases and therefore we investigated the roles of annotated proteins known to affect 3’ to 5’ RNA degradation. We learned that mutations of *REX2*, *REX3*, or *REX4*, and *POP2*, did not affect levels of any of the 8 tested families of tRNA introns (Supplementary Table 3). Concerning the exosome, mutation of *RRP6*, encoding the nucleus-located catalytic RNase of the exosome, also did not affect the abundance of any of the 8 tested tRNA introns. On the other hand, cells bearing a mutation of *DIS3*, specifically the *dis3-1 and dis3-ts* alleles, encoding the other exosome catalytic RNase, accumulated elevated levels of the tRNA^Leu^_CAA_ and tRNA^Phe^_GAA_ introns, but curiously only at the permissive temperature (Fig. 4; Supplementary Table 4; Supplemental Fig. 3B); since the tRNA^Leu^_CAA_ intron levels are Xrn1-dependent, this intron may be destroyed bidirectionally. Surprisingly, none of the numerous other exosome subunits or associated factors tested positive for tRNA intron accumulation. The results indicate that: (1) 3’ to 5’ turnover doesn’t appear to be a major mechanism for tRNA intron turnover as only tRNA^Leu^_CAA_ and tRNA^Phe^_GAA_ intron turnover are affected by Dis3; (2) tRNA^Leu^_CAA_ intron turnover appears to be bidirectional; (3) the tRNA^Ser^_GCU_ intron seems not to be destroyed by either 5’ to 3’ or 3’ to 5’ nucleases (Fig. 4). However, a caveat of these conclusions is that we scored only yeast with single exosome mutations and therefore we cannot eliminate roles for proteins which function redundantly in 3’ to 5’ RNA turnover.

#### Endonucleolytic tRNA intron turnover

We identified three endonucleases which impact the levels of tRNA introns. Mutations of *LAS1*, *NOB1,* and *YSH1* each resulted in elevated levels of subsets of tRNA introns. The levels of each of the tRNA intron families were affected by at least one of the tested endonucleases and most of the tested tRNA intron families were affected by different subsets of endonucleases (Supplementary Table 4; Supplementary Fig. 3C). Thus, despite the fact that all detected tRNA introns appear to be full length (Fig. 1), endonucleolytic activity appears to play a major role in the turnover of free tRNA introns as detailed below and diagramed in Fig. 4).

Las1: Mutation of *LAS1* resulted in accumulation of 5 tRNA introns (tRNA^Ile^_UAU_, tRNA^Leu^_CAA_, tRNA^Trp^_CCA_, tRNA^Lys^_UUU_, and tRNA^Ser^_GCU_) even though as predicted by the RNAfold web server (http://rna.tbi.univie.ac.at/cgi-bin/RNAWebSuite/RNAfold.cgi) none of these intron sequences is predicted to have the previously described Las1 recognition site described for pre-rRNA processing which is comprised of a stem with an unpaired A followed 2 paired G-Cs (45). Las1 interacts with Grc3 which enhances Las1’s activity (45, 46). Las1 cleavage generates upstream oligonucleotides bearing 2’, 3’ cyclic phosphates and downstream oligonucleotides at the 3’ end of the cut site bearing a 5’ hydroxyl which could be phosphorylated by the Grc3 5’ kinase (45) (or Trl1?), thereby possibly leading to subsequent additional internal destruction by Xrn1 exonucleolytic turnover of the tRNA^Ile^_UAU_ and/or tRNA^Leu^_CAA_ introns which accumulate in *grc3 or trl1, las1,* and *xrn1* mutant yeast (Fig. 4, dotted arrows).

Nob1: tRNA^Ile^_UAU_ and tRNA^Pro^_UGG_ introns accumulate in the verified *nob1-ts* mutant upon 2 hr incubation at the nonpermissive temperature (Supplemental Fig. 3C). Nob1 is a Pin domain containing RNase-like endonuclease functioning in pre-rRNA processing. Nob1 cleavage generates 3’ hydroxyl and 5’ P-containing fragments. Nob1 cuts 20S pre-rRNA at the single stranded D site to generate mature 18S rRNA (37). Since tRNA^Pro^_UGG_ intron accumulates in both *nob1-ts* and *xrn1* mutant cells it is possible that upon endonucleolytic cleavage by Nob1 the 5’ phosphorylated fragment is subsequently further degraded by Xrn1 5’ to 3’ exonuclease (Fig. 4).

Ysh1: Ysh1 is an essential endoribonuclease which is a component of the cleavage and polyadenylaton factor functioning in newly transcribed mRNA 3’ end processing; it has also been reported to function in snoRNA processing and mRNA splicing (36). Upon incubation WT and *ysh1-ts* cells at the *ysh1* nonpermissive temperature tRNA^Ile^_UAU_, tRNA^Leu^_CAA_, tRNA^Pro^_UGG_, tRNA^Lys^_UUU_, and tRNA^Ser^_GCU_, but not tRNA^Trp^_CCA_, tRNA^Phe^_GAA_ or tRNA^Ser^_CGA_ introns accumulate in *ysh1-ts* cells relative to WT and accumulation is complemented by a low copy vector encoding WT *YSH1* (Supplemental Fig. 3C). Accumulation of the tRNA introns could result from direct endonucleolytic cleavage of the introns (Fig. 4) or, alternatively, given Ysh1’s well described function in mRNA processing/polyadenylation, tRNA intron accumulation in *ysh1-ts* mutant cells may result indirectly from downregulation of mRNAs.

#### Roles for nucleases involved in the cellular levels of circular tRNA^Trp^_CCA_ intron

Given that Las1 endonuclease appears to play a role in the cellular levels of linear tRNA^Trp^_CCA_ intron, it stands to reason that turnover of the circular form could utilize the same pathways following initial Las1-mediated linearization. However, degradation of the linear and circular forms of the tRNA^Trp^_CCA_ intron utilizes only partially overlapping pathways. For example, elevated levels of the linear and circular forms of the tRNA^Trp^_CCA_ intron accumulate in *xrn1Δ* cells, indicating similar 5’ to 3’ turnover. However, the pathways of endonucleolytic digestion differ. The linear, but not circular, tRNA^Trp^_CCA_ intron accumulates in *las1-ts* cells at the non-permissive temperature, whereas the circular, but not the linear, tRNA^Trp^_CCA_ intron accumulates in *nob1-ts* cells at the non-permissive temperature. Linearization of the tRNA^Trp^_CCA_ intron by Nob1 could render it susceptible to 5’ to 3’ degradation by Xrn1 (Fig. 4)

#### Summary of studies of annotated enzymes involved in RNA turnover

The results of our candidate screen for roles of annotated RNases and kinases in the cellular levels of 8 of the 10 families of tRNA introns document that tRNA intron levels are impacted by subsets of endo- and exonucleases. Surprisingly, most of the tRNA intron families surveyed are affected by different subsets of tested enzymes (Fig. 4; Supplementary Table 4; Supplemental. Fig. 3). Given the known roles of these enzymes in RNA turnover, many of the effects on tRNA intron levels are likely direct, however, indirect effects cannot be ruled out.

### Growth/environmental conditions impacting tRNA intron levels

We previously provided evidence that free tRNA introns (fitRNAs) function as small noncoding RNAs that contribute to both basal and stress-triggered post-transcriptional gene expression of target mRNAs via complementarity of the target mRNAs with the fitRNAs and subsequent mRNA degradation (17). However, our studies were restricted to just two families of tRNA introns, tRNA^Ile^_UAU_ and tRNA^Trp^_CCA_, and for cells in early log phase or cells exposed to hydrogen peroxide (H_2_O_2_) stress. To obtain a more thorough understanding of the roles of all families of fitRNAs in target mRNA levels under both normal and stress conditions we expanded our studies to assess the levels of free tRNA introns for cells grown in both rich (YEPD) and synthetic complete (SCD) media from early log through stationary phase as well as for cells exposed to a variety of stress conditions. Cell growth was monitored by OD_600_ as well as by cell number throughout the growth cycle with hourly time points (Supplementary Fig. 4). RNAs were extracted from the cells for each time point, followed by northern blot analyses of the levels of each of 8 of the10 tRNA intron families employing tRNA intron-specific probes (Fig. 5). For the tRNA^Trp^_CCA_ intron we assessed the levels of both the linear and circular forms. As the rate of tRNA transcription and processing varies throughout the growth curve (Fig. 5; Supplementary Fig. 5), we assessed the relative levels of tRNA introns as the ratio of tRNA introns to their pre-tRNA counterparts and normalized to levels of 5S rRNA.

**Figure 5.**
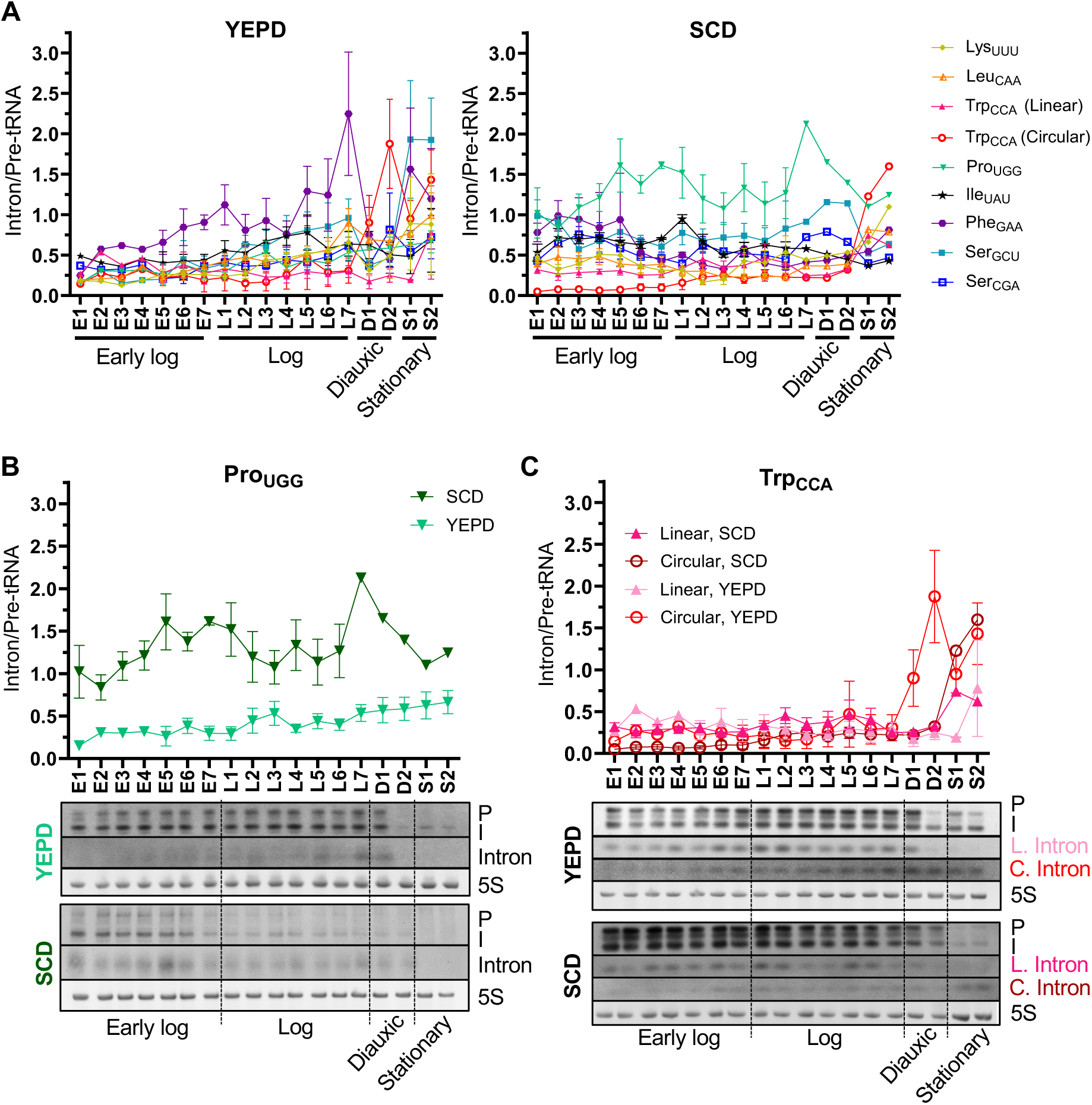
tRNA introns accumulate at varied levels in different media and in different phases of growth curves. **A.** Quantitation of free intron levels for the different intron-containing tRNA families in cells grown in YEPD (left) or SCD (right) media. Cells were inoculated in early log phase and samples for RNA extraction were taken hourly from early log phase to stationary phase. Early log phase (E1-7), log phase (L1-7), diauxic shift (D1-2), and stationary phase (S1-2) were determined by the amount of cells and position of the growth curve (Supplementary Fig. 4). RNAs were extracted and subject to northern blot analyses. Blots were probed with oligonucleotides complementary to the indicated tRNA intron (also see Supplementary Fig. 5 for Northern blots). Intron levels are quantified as the free intron levels/pre-tRNA levels, relative to 5S rRNA levels. **B, C.** Northern blots and quantification of tRNA^Pro^_UGG_ (B) and tRNA^Trp^_CCA_ (C) intron levels for WT cells grown in YEPD vs. SCD media, as in (A). P: initial pre-tRNA transcripts; I: intron-containing end-processed pre-tRNA;

The results provided unanticipated insights. First, the levels of individual tRNA introns differed in glucose-containing yeast extract peptone (YEPD) vs. synthetic complete (SCD) media (Fig. 5; Supplementary Fig. 5). For example, the intron/pre-tRNA levels for the tRNA^Phe^_GAA_ intron was highest among the tRNA introns families throughout the YEPD media growth curve, but relatively the same as the majority of the other tRNA intron families in cells grown in SCD (Supplementary Fig. 5). Conversely, levels of the tRNA^Pro^_UGG_ intron were low throughout the YEPD growth curve but the highest among the tRNA intron families for cells grown in SCD media (Fig. 5A and B). Second, in both types of media the linear tRNA^Trp^_CCA_ intron was expressed at similar intron/pre-tRNA levels from early log to stationary phase. In contrast, the circular tRNA^Trp^_CCA_ intron changed significantly as it was at barely detectable levels from early log to late log phase, but at significantly elevated levels at the diauxic shift and stationary phase in YEPD media and most elevated at stationary phase for SCD media (Fig. 5A and C). Since we previously reported that fitRNAs negatively regulate subsets of mRNAs that bear complementarity to individual tRNA introns (17), family-specific differences in levels of fitRNAs throughout the growth cycle and in different media seem likely to impact the subsets of mRNAs bearing complementarity to individual introns and, hence the proteome. Future studies will test this prediction.

### Environmental stress impacts tRNA intron levels

Exposure of yeast to hydrogen peroxide (H_2_O_2_) stress resulted in increased levels of tRNA^Trp^_CCA_ and tRNA^Leu^_CAA_ introns but did not affect the levels of the other tested tRNA introns (17). To obtain a fuller appreciation of the relationship of tRNA intron levels with respect to cellular stresses, we studied levels of tRNA introns upon other stress conditions.

We first explored whether, like H_2_O_2_, oxidative stress induced by exposure to diamide or dithiotreitol (DTT) also results in increased levels of tRNA introns. H_2_O_2_ levels varying from 0.5 mM (17) to 3.0 mM (Supplementary Fig. 6) resulted in tRNA transcription termination within 30 min and concomitant elevated levels of the tRNA^Trp^_CCA_ or tRNA^Leu^_CAA_ introns. In contrast, varying diamide levels from 1.5 to 5 mM [1.5 mM exposure affects mRNA levels (47)] did not result in tRNA^Trp^_CCA_ nor tRNA^Leu^_CAA_ intron accumulation, even though tRNA transcription ceased at 5 mM treatment. Levels of tRNA transcription were not reduced by exposure to DTT ranging from 2.5 mM to 10mM, [2.5 mM exposure affects mRNA levels (47)], although the tRNA^Trp^_CCA_ intron but not the tRNA^Leu^_CAA_ intron showed slight accumulation at 10 mM treatment after 60 min exposure (Supplementary Fig. 6). Thus, not all inducers of oxidative stress result in tRNA intron accumulation.

We also tested other types of environmental stress including amino acid or nitrogen deprivation and growth in media with alternative carbon sources including galactose, raffinose, or ethanol. No significant effects upon tRNA intron levels were observed (Supplementary Figs. 7, 8, 9). In contrast, exposure of cells to heat stress at 42°C resulted in elevated levels of a single tRNA intron family, tRNA^Pro^_UGG_. After 30 min at 42°C, tRNA^Pro^_UGG_ levels increased 1.5-fold, reaching >2-fold by 60 min and elevated levels were maintained for at least 2 hr. (Fig. 6; Supplementary Fig. 10). tRNA^Pro^_UGG_ is encoded by 10 genes bearing 4 slightly different intron sequences ranging from 30-33 nucleotides (1). The tRNA^Pro^_UGG_ introns accumulating under this heat stress migrate slower than those detected when cells are grown at 23°C (Fig. 6), perhaps indicative of accumulation of the longer subset of the encoded tRNA^Pro^_UGG_ introns being affected by this heat stress. Changes in the cellular levels of tRNA intron families may function in regulation of the proteome in response to environmental stress.

**Figure 6.**
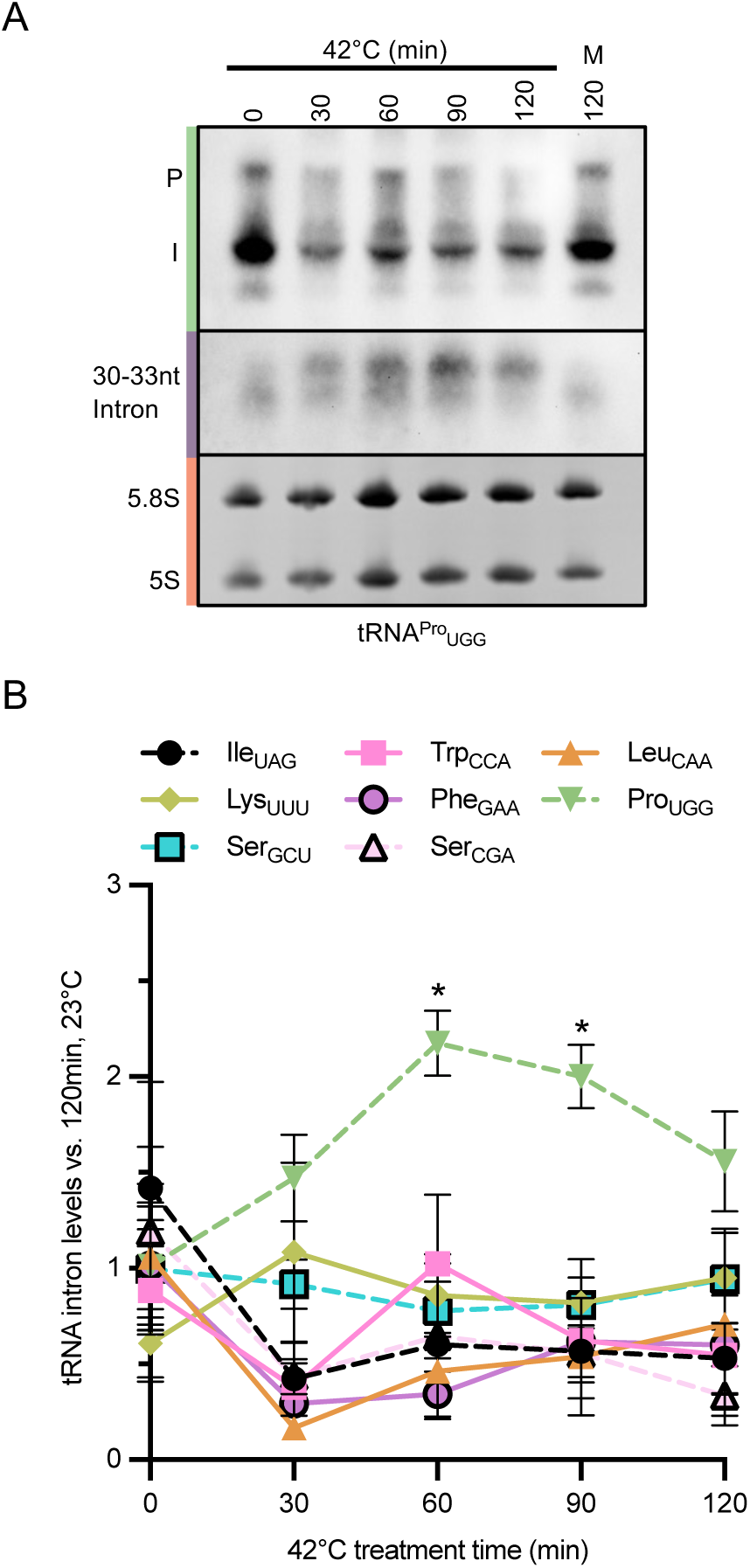
Heat stress causes increased tRNA^Pro^_UGG_ intron levels. **A.** WT cells were grown 23^0^C to early log phase and then were shifted to 42°C for 0-120 min. Mock treated cells (M) were harvested at the same time as 2 hr.-treated cells. Small RNAs were isolated and northern blot analysis was performed using oligonucleotides complementary to the tRNA^Pro^_UGG_ intron. Representative northern blot showing levels of precursor tRNA^Pro^_UGG_ (P, initial transcript; I, intron-containing end-processed pre-tRNA [green bar]) or free intron (purple bar). 5.8S and 5S rRNA (orange bar) serve as loading controls. **B.** Quantitation of tRNA intron levels from the different intron-containing tRNA families, as determined by the northern blot analysis (see Supplementary Fig. 10). Data are expressed relative to untreated cells, set as 1. n = 3. Data are expressed as mean ± SEM. \**p* < 0.05.

## DISCUSSION

The presence of introns within a subset of tRNA genes is nearly ubiquitous throughout archaea and eukaryotes. Despite the fact that tRNA introns from different tRNA families have distinct sequences and lengths, and consequently pre-tRNA anticodon loop structures, all families of intron-containing tRNAs are spliced by a single, conserved mechanism using the SEN complex via subunit interactions with the mature tRNA tertiary structure (3). Spliced, mature tRNAs are one of the most long-lived RNAs in the cell (48), with repair mechanisms such as CCA 3’ addition (49), in place to repair those damaged tRNAs, highlighting their essential constitutive role in the cell. In contrast, the freed tRNA introns are very short-lived and barely detectable under standard growth conditions. However, as our recent published studies (17) and data provided here demonstrate, the importance of tRNA introns is evident not by their longevity but rather by the numerous and distinct mechanisms to control their levels in tRNA family-specific, stress-specific manners in order to meet the ever-changing needs of the cell.

The surprising discovery that all tRNA introns are not degraded by the same two-step process involving 5’ phosphorylation by Trl1 and subsequent 5’ to 3’ turnover by Xrn1, as had been previously identified for tRNA^Ile^_UAU_ (15) led us to screen for other annotated yeast RNA kinases and 5’ to 3’ nucleases that may substitute for these enzymes. Although we identified a role for Grc3, the only other known RNA kinase, in the turnover of two tRNA intron families, these were not introns whose turnover was Trl1- or Xrn1-independent, respectively. Specifically, none of the 3 intron families whose turnover is independent of Trl1, tRNA^Trp^_CCA_, tRNA^Pro^_UGG_ and tRNA^Ser^_CGA_, displayed intron accumulation in a Grc3-defective strain. Of the introns whose turnover was independent of Xrn1, specifically tRNA^Lys^_UUU_, tRNA^Phe^_GAA_, tRNA^Ser^_GCU_ and tRNA^Ser^_CGA_, none displayed intron accumulation in cells deficient in any of the other 5’ to 3’ exonucleases, Dxo1 or Rat1. Thus, it appears that not all tRNA intron machinery has been identified.

Instead of identifying the missing components involved in intron turnover, our screen of annotated nucleases affecting tRNA intron cellular levels identified novel components and expanded our understanding of the multitude of mechanisms at play. For example, the tRNA^Ile^_UAU_ intron, which is degraded by a Trl1/Xrn1-mediated mechanism is also subject to cleavage by two endonucleases. The turnover of the tRNA^Leu^_CAA_ intron, which is also Trl1 and Xrn1-dependent, utilizes an additional RNA kinase, Grc3, and a different combination of endonucleases. Further, turnover of the tRNA^Leu^_CAA_ intron is dependent on Dis3, indicating that the tRNA^Leu^_CAA_ intron is degraded via 5’ to 3’, 3’ to 5’, and endonucleolytic activities. In accordance with these findings, differing combination of turnover components were identified for nearly all of the 8 introns assessed (Fig. 4; Supplementary Table 4). Given the multitude of turnover pathways for a given tRNA intron family and the barely detectable levels under standard conditions, it is remarkable that loss of a single component is sufficient to lead to appreciable intron accumulation. For example, the tRNA^Leu^_CAA_ intron accumulates in *xrn111* cells, and thereby is degraded by the Xrn1 5’ to 3’ nuclease, and also in temperature sensitive *dis3* cells grown at the non-permissive temperature, which functions in 3’ to 5’ turnover. The data suggest that these pathways may not simply be redundant, but they may have distinct roles in regulating levels of tRNA introns.

Despite the information gained by investigations of annotated nucleases several questions remain. First, information is lacking regarding 3’ to 5’ turnover of tRNA introns. Of the 8 introns tested, Dis3 mutations resulted in accumulation of tRNA^Leu^_CAA_ and tRNA^Phe^_GAA_, but none of the other exosome subunits tested affected the levels of tRNA introns. Possibly exosome subunit overlapping functions obscure the phenotypes of ablation of single genes. Secondly, it is unclear why the tRNA^Trp^_CCA_ intron circularizes whereas the other tested tRNA introns accumulate as linear molecules, when all tRNA introns possess 5’ hydroxyl and 2’, 3’ cyclic phosphates upon cleavage by the SEN complex. Recently, the enzyme-RNA substrate binding determinants for the *Chaetomium thermophilum* Trl1 ligase were described; they contain sequence nonspecific 4-6 nt stem-loop structures with 5’ and 3’ overhangs (50). As predicted by the RNAfold web server (http://rna.tbi.univie.ac.at/cgi-bin/RNAWebSuite/RNAfold.cgi), the *S. cerevisiae* tRNA^Trp^_CCA_ intron is predicted to possess a stem-loop structure with 5’ and 3’ overhangs, but so do other tRNA introns that do not form circles *in vivo*. Furthermore, circularization does not appear to be simply a consequence of the amount of linear intron, since *las1-ts* cells at the non-permissive temperature accumulate linear but not circular tRNA^Trp^_CCA_ intron (Supplementary Fig. 3C). Conversely, *nob1-ts* cells at the non-permissive temperature have more circular but not linear tRNA^Trp^_CCA_ intron (Supplementary Fig. 3). Additionally, WT yeast cells grown in YEPD and SCD display increases in circular tRNA^Trp^_CCA_ intron levels as cells enter stationary phase, despite little change in the linear form, relative to precursor tRNA^Trp^_CCA_ levels (Fig. 5). Thus, the specificity of tRNA^Trp^_CCA_ circularization by Trl1 remains an enigma. Third, one might have predicted that only Trl1 5’ kinase activity would be required to render tRNA introns for turnover by 5’ to 3’ exonucleases. However, we learned that the tRNA^Ile^_UAU_ intron levels are altered by mutation of all three enzyme domains; moreover, turnover of the remaining Trl1-dependent introns only requires Trl1 kinase and CPD activities (Fig. 3). Finally, despite considerable effort, we failed to understand why only the tRNA^Leu^_CAA_ intron smears towards a slower migration when Trl1 kinase activity is defective. Future studies of the structure/sequences of this tRNA intron in Trl1 kinase defective cells should uncover the reason for aberrant migration.

We learned that growth conditions and environmental stresses alter family-specific tRNA introns cellular levels. We anticipated that under standard laboratory growth conditions tRNA intron levels would be solely dependent upon tRNA transcription and processing. However, this is not the case as the ratios of tRNA introns/pre-tRNAs varied between YEPD and SCD media and differed among the tRNA families. Further, tRNA intron levels changed throughout the growth curves; this was particularly prominent for the tRNA^Trp^_CCA_ circular intron which accumulated to its highest levels at the diauxic shift/stationary phase. We had also anticipated that the levels of tRNA introns might be altered upon nutrient deprivation via induction of the integrated stress response; surprisingly, we did not detect tRNA intron level changes in response to amino acid or nitrogen deprivation or by changes in carbon source. We also extended studies of the tRNA intron levels in response to various environmental stresses. Regarding oxidative stress we previously reported that tRNA^Trp^_CCA_ and tRNA^Leu^_CAA_ introns accumulate upon H_2_O_2_ exposure (17). Here we tested oxidative stresses by DTT and diamide and learned that these reagents did not cause changes in tRNA intron levels. The results are not surprising as the regulatory networks for H_2_O_2_ vs DTT vs diamide differ (47). Regarding heat stress, we uncovered a tRNA family-specific response to heat shock for tRNA^Pro^_UGG_ upon exposure to 42°C. Thus, among the stresses tested, oxidative and heat stress result in increased levels of different subsets of tRNA introns. It is quite possible that different turnover pathways could be preferentially used under different stress conditions. Future studies of other environmental stresses will provide a fuller appreciation of the links of tRNA intron turnover and environmental stress pathways.

It will be important to delineate the subcellular location(s) of tRNA intron turnover. Our studies documenting 5’ to 3’ turnover by cytoplasmic Xrn1 but not nuclear Rat1 support the model that tRNA intron turnover is a cytoplasmic process [(15) and these data]. Likewise, Nob1 is a cytoplasmic endonuclease (37, 51). However, several of the kinases/nucleases that affect tRNA intron levels are not cytoplasmic proteins. RNA kinase Grc3 and endonuclease Las1 are reported to be nuclear residents (51–53). It is unknown whether there are undetected cytoplasmic pools of these proteins or whether, like for full length tRNAs, free tRNA introns are able to transit in a retrograde fashion from the cytoplasm to the nucleus (or other subcellular locations) after splicing at the cytoplasmic mitochondrial surface (54, 55). Future imaging of the subcellular locations of individual free introns under both normal and stress conditions will be informative.

Together, these findings highlight the many cellular pathways utilized to regulate tRNA intron levels and the specificity of these pathways for different tRNA families and varying cellular conditions. Surprisingly, of all the different classes of nucleases studied, endonucleases appear to play a significant role in tRNA intron turnover, as this class of nucleases are involved in the degradation of 6 of the 8 tRNA intron families studied. Given our previous findings identifying the tRNA^Ile^UAU and tRNA^Trp^_CCA_ introns as negative regulators of gene expression via binding to ORFs of target mRNAs bearing complementarity (17), the role of endonucleases is intriguing, as different endonucleases could cleave tRNA introns at different sites, generating “mini-fitRNAs” that could regulate the gene expression of distinct sets of transcripts. Conversely, it is possible that the circular tRNA^Trp^_CCA_ could function as an additional fitRNA via its junction of 5’ and 3’ nucleotides. Overall, the full range of tRNA intron functions in yeast remains to be discovered. Given their evolutionary conservation, it is likely that tRNA introns also play important roles in human health and disease.

## METHODS

### Yeast strains and growth

The yeast strains employed, and their sources are listed in Supplementary Table 2. For most experiments, yeast strains were propagated in rich YEPD (yeast extract with peptone and 2% glucose) media at 23° or 30°C to early log phase. For studies employing complementation by genes harbored on plasmids or for the studies of the roles of Trl1 domains in tRNA intron levels, the yeast strains were grown in synthetic complete glucose containing media (SCD) lacking the appropriate component for selection of the plasmids. To analyze phenotypes of cells with mutations of essential genes, the strains were propagated to early log phase at 23°C and then shifted to 37°C for 2 hr before harvesting.

For studies of tRNA introns during various growth phases cells were grown overnight in either YEPD or SCD to early log phase. For each sample of log phase growth, ranging from 0 to 15 hr, the cell densities were determined by OD_600_ and by hemocytometer (to determine cells/mL) prior to harvesting. The cultures were grown overnight to obtain cells in stationary phase.

### Environmental stress conditions

Cells were grown under the conditions indicated for each specific stress condition, then harvested by centrifugation at 3,000 x g for 5 min at 4°C to pellet the cells. Cell pellets were washed one time in the same media they were grown in, then stored at -80°C.

Heat stress – Cells were grown to early log phase at 23°C and then shifted to 42°C for the indicated time from 0 to 120 min. prior to harvesting. Mock cultures remained at 23^0^C for 120 min.

Oxidative stress – Cells were grown at 23^0^C to OD_600_ in YEPD media. H_2_O_2_, diamide, or DTT were added to the indicated concentrations and samples were harvested starting at 10 sec. and at each of the indicated time points. Mock cultures lacking H_2_O_2_, diamide, or DTT were harvested at the indicated times.

Nutrient deprivation - Cells were grown in SCD media at 23°C to OD_600_ of 0.3 – 0.4. Samples were harvested at the indicated times.

Alternative carbon sources – Cells were grown overnight in YP (yeast extract with peptone) and 2% of one of the following carbon sources: glucose, galactose, raffinose, or EtOH. Cells were grown to an OD_600_ of 0.3 – 0.4, then harvested at the indicated times. The OD_600_ at the time of harvesting is also indicated, since the growth of yeast cells is greatly impacted by the carbon source.

### Screen for gene products involved in tRNA intron turnover and verification analyses

For the tRNA intron turnover screen, gene products were based on their annotated roles in any step of RNA turnover. The 32 genes (Supplementary Table 3) included RNA kinases, exoribonucleases, and endoribonucleases that were available in yeast commercially-available mutant deletion collections (Open Biosystems) and two temperature sensitive collections provided by Drs. Boone and Heiter (20, 24). Levels of tRNA introns from each strain were assessed (see below) at least 2 times and mutants for which at least 1 of the 8 analyzed tRNA introns levels increased were further confirmed. Since the mutant collections contain errors (56), the phenotypes of each lesion was verified by assessing tRNA intron levels from independently derived strains, by complementation employing plasmids from the molecular barcoded yeast ORF library (MOBY) bearing the WT counterpart of the mutant genes (57, 58) and/or by RNA analyses from cells grown at permissive vs. nonpermissive temperatures. The *nob1-ts* mutation was verified by assessment of its published defect in processing pre-rRNA (59). The results of the screen are summarized in Fig. 4, Supplementary Table 4.

### RNA isolation

“Small” RNAs, including 5S and 5.8S rRNAs, tRNAs, and tRNA introns, were obtained from cells grown to early log phase (OD_600_ 0.3), unless otherwise indicated, by whole-cell phenol extraction as previously described (15, 19). RNAs from temperature sensitive mutants were obtained following incubation at the nonpermissive temperature (37°C) for 2 hr.

### TEX treatment of RNA

Ten micrograms of small RNAs were dissolved in nuclease-free water and incubated with 1 unit of terminator exonuclease (TEX; Lucigen) for 1 hr at 30°C according to the manufacturer’s protocol. The reaction was terminated by adding 1μL of 100mM EDTA (pH 8.0). RNA was subsequently used for northern blot analysis.

### Northern blot analyses

RNAs were resolved in 10% polyacrylamide, 8 M urea gels and transferred to Hybond-N+ nylon membranes (Amersham) as previously detailed (15, 19). Following transfer, the RNAs were UV crosslinked to the membrane. Digoxigenin (DIG)-tagged DNA oligonucleotides complementary to tRNA intron sequences or the tRNA^Leu^_CAA_ 5’ exon (Supplementary Table 1) were hybridized to the membrane, followed by anti-DIG antibody conjugated with alkaline phosphatase. Membranes were incubated with CDP-Star (Roche) chemiluminescent substrate and chemiluminescent signals were detected by using a UVP ChemStudio instrument from Analytic Jena. Quantification of the intensity of the bands was achieved using Image J software.

### Genomic DNA isolation and DNA sequencing

To determine the DNA sequence of the *trl1-ts* allele, genomic DNA was first isolated by transferring 100-200μL of *trl1-ts* yeast culture (OD_600_ = 0.2-0.6) to a microcentrifuge tube and centrifuging at 15,000 x g for 1 min at room temperature to pellet cells. The supernatant was removed and cells were resuspended in 100μL of 200mM lithium acetate with 1% SDS. The cell suspension was incubated at 70°C for 15 min. Following incubation, 300μL of 96% ethanol was added and samples were mixed by brief vortexing. DNA was collected by centrifugation at 15,000 x g for 3 min at room temperature. The DNA was washed with 100μL of 70% ethanol, then air dried at room temperature for 5-10 min. DNA was dissolved in 100μL of TE buffer [10mM Tris (pH 8.0), 1mM EDTA]. Cell debris was pelleted by centrifugation at 15,000 x g for 1 min at room temperature. Amplification of the trl1-ts allele was conducted by PCR using 1-3 *μL* (∼500-1000ng/*uL*) of genomic DNA, the forward and reverse primers SM62 and SM63, respectively (Supplementary Table 1), and the Q5 High-Fidelity DNA polymerase (NEB). The PCR product was visualized by gel electrophoresis on an ethidium bromide-stained 1% agarose gel. The PCR product was cut out and purified using the QIAquick Gel extraction kit (Qiagen). Sequencing, using the same primers as for PCR, was performed by Plasmidsaurus using Oxford Nanopore Technology with custom analysis and annotation.

### Quantification and Statistical Analyses

All data are expressed as mean ± SEM. N values indicate the number of biological replicates per experiment. Each experiment was performed at least in triplicate (unless otherwise indicated) with the results from one representative Northern blotting experiment displayed in each figure. Statistical analyses were performed using the GraphPad Prism 9 software. One-way ANOVA with Dunnett’s multiple comparisons test was used when comparing more than two groups. A p value of <0.05 was considered significant. Statistical significance is denoted in all figures as: * p <0.05; ** p<0.01; *** p<0.001; **** p<0.0001; ns p>0.05.

## Supporting information

Metcalf et al Suppl. Materials

## ACKNOWLEGEMENTS

This work was supported by funding from the National Institutes of Health, grant number GM122884 to A.K.H., Pelotonia Undergraduate Fellowships to P.L.S. and A.B. and Undergraduate Research Scholarships (URS) to P.L.S. and KC. We thank Drs. Beate Schwer and Steward Shulman for yeast strains and plasmids and Dr. Ambro van Hoof for advice. We thank Drs. C. Boone and P. Heiter for providing the yeast temperature sensitive mutant collections. We also thank all members of the Hopper lab for scientific conversations and comments on the manuscript.

## AUTHOR CONTRIBUTIONS

Conceptualization: SM, RTN, AB, and AKH; Investigation: SM, RTN, AB, KC, and PLS; Writing: SM, RTN, PLS, and AKH; Supervision: AKH.

## REFERENCES

1. P. P. Chan, T. M. Lowe, GtRNAdb 2.0: an expanded database of transfer RNA genes identified in complete and draft genomes. Nucleic Acids Res 44, D184–189 (2016).

2. A. van Hoof, T. Furata, S. Arur, A eukaryote without tRNA introns. RNA 10.1261/rna.080669.125 (2025).

3. C. K. Hayne et al., Structural basis for pre-tRNA recognition and processing by the human tRNA splicing endonuclease complex. Nat Struct Mol Biol 30, 824–833 (2023).

4. E. M. Phizicky, A. K. Hopper, The life and times of a tRNA. RNA 29, 898–957 (2023).

5. G. Knapp, R. C. Ogden, C. L. Peebles, J. Abelson, Splicing of yeast tRNA precursors: structure of the reaction intermediates. Cell 18, 37–45 (1979).

6. C. A. Schmidt, A. G. Matera, tRNA introns: Presence, processing, and purpose. Wiley Interdiscip Rev RNA 11, e1583 (2020).

7. S. V. Paushkin, M. Patel, B. S. Furia, S. W. Peltz, C. R. Trotta, Identification of a human endonuclease complex reveals a link between tRNA splicing and pre-mRNA 3’ end formation. Cell 117, 311–321 (2004).

8. T. Yoshihisa, Handling tRNA introns, archaeal way and eukaryotic way. Front Genet 5, 213 (2014).

9. S. Shuman, RNA Repair: Hiding in Plain Sight. Annu Rev Genet 57, 461–489 (2023).

10. E. M. Phizicky, S. A. Consaul, K. W. Nehrke, J. Abelson, Yeast tRNA ligase mutants are nonviable and accumulate tRNA splicing intermediates. J Biol Chem 267, 4577–4582 (1992).

11. G. M. Culver, S. M. McCraith, S. A. Consaul, D. R. Stanford, E. M. Phizicky, A 2’-phosphotransferase implicated in tRNA splicing is essential in Saccharomyces cerevisiae. J Biol Chem 272, 13203–13210 (1997).

12. S. Ghosh, G. Wimberly-Gard, A. Jacewicz, B. Schwer, S. Shuman, Identification, characterization, and structure of a tRNA splicing enzyme RNA 5’-OH kinase from the pathogenic fungi Mucorales. RNA 30, 1674–1685 (2024).

13. K. S. Ahammed, A. van Hoof, Fungi of the order Mucorales express a “sealing-only” tRNA ligase. RNA 30, 354–366 (2024).

14. C. A. Schmidt, J. D. Giusto, A. Bao, A. K. Hopper, A. G. Matera, Molecular determinants of metazoan tricRNA biogenesis. Nucleic Acids Res 47, 6452–6465 (2019).

15. J. Wu, A. K. Hopper, Healing for destruction: tRNA intron degradation in yeast is a two-step cytoplasmic process catalyzed by tRNA ligase Rlg1 and 5’-to-3’ exonuclease Xrn1. Genes Dev 28, 1556–1561 (2014).

16. Z. Lu et al., Metazoan tRNA introns generate stable circular RNAs in vivo. RNA 21, 1554–1565 (2015).

17. R. T. Nostramo et al., Free introns of tRNAs as complementarity-dependent regulators of gene expression. Mol Cell 85, 726–741 e726 (2025).

18. A. Stevens, Purification and characterization of a Saccharomyces cerevisiae exoribonuclease which yields 5’-mononucleotides by a 5’ leads to 3’ mode of hydrolysis. J Biol Chem 255, 3080–3085 (1980).

19. J. Wu, H. Y. Huang, A. K. Hopper, A rapid and sensitive non-radioactive method applicable for genome-wide analysis of Saccharomyces cerevisiae genes involved in small RNA biology. Yeast 30, 119–128 (2013).

20. S. Ben-Aroya et al., Toward a comprehensive temperature-sensitive mutant repository of the essential genes of Saccharomyces cerevisiae. Mol Cell 30, 248–258 (2008).

21. K. L. Patrick et al., Distinct and overlapping roles for two Dicer-like proteins in the RNA interference pathways of the ancient eukaryote Trypanosoma brucei. Proc Natl Acad Sci U S A 106, 17933–17938 (2009).

22. B. Schwer, A. Aronova, A. Ramirez, P. Braun, S. Shuman, Mammalian 2’,3’ cyclic nucleotide phosphodiesterase (CNP) can function as a tRNA splicing enzyme in vivo. RNA 14, 204–210 (2008).

23. E. A. Winzeler et al., Functional characterization of the S. cerevisiae genome by gene deletion and parallel analysis. Science 285, 901–906 (1999).

24. Z. Li et al., Systematic exploration of essential yeast gene function with temperature-sensitive mutants. Nat Biotechnol 29, 361–367 (2011).

25. P. Braglia, K. Heindl, A. Schleiffer, J. Martinez, N. J. Proudfoot, Role of the RNA/DNA kinase Grc3 in transcription termination by RNA polymerase I. EMBO Rep 11, 758–764 (2010).

26. C. K. Hayne, C. A. Schmidt, M. I. Haque, A. G. Matera, R. E. Stanley, Reconstitution of the human tRNA splicing endonuclease complex: insight into the regulation of pre-tRNA cleavage. Nucleic Acids Res 48, 7609–7622 (2020).

27. Y. Xue et al., Saccharomyces cerevisiae RAI1 (YGL246c) is homologous to human DOM3Z and encodes a protein that binds the nuclear exoribonuclease Rat1p. Mol Cell Biol 20, 4006–4015 (2000).

28. J. H. Chang et al., Dxo1 is a new type of eukaryotic enzyme with both decapping and 5’-3’ exoribonuclease activity. Nat Struct Mol Biol 19, 1011–1017 (2012).

29. X. Ye, A. Axhemi, E. Jankowsky, Alternative RNA degradation pathways by the exonuclease Pop2p from Saccharomyces cerevisiae. RNA 27, 465–476 (2021).

30. A. Keidel, C. L. Long, J. Iwasa, E. Conti, RNA-Degrading Exosome Complexes: Molecular Mechanisms and Structural Insights. Annu Rev Cell Dev Biol 10.1146/annurev-cellbio-111822-115115 (2025).

31. K. Schmidt, J. S. Butler, Nuclear RNA surveillance: role of TRAMP in controlling exosome specificity. Wiley Interdiscip Rev RNA 4, 217–231 (2013).

32. C. Sidrauski, P. Walter, The transmembrane kinase Ire1p is a site-specific endonuclease that initiates mRNA splicing in the unfolded protein response. Cell 90, 1031–1039 (1997).

33. D. M. Thompson, R. Parker, The RNase Rny1p cleaves tRNAs and promotes cell death during oxidative stress in Saccharomyces cerevisiae. J Cell Biol 185, 43–50 (2009).

34. L. Gasse, D. Flemming, E. Hurt, Coordinated Ribosomal ITS2 RNA Processing by the Las1 Complex Integrating Endonuclease, Polynucleotide Kinase, and Exonuclease Activities. Mol Cell 60, 808–815 (2015).

35. S. M. Barabino, M. Ohnacker, W. Keller, Distinct roles of two Yth1p domains in 3’-end cleavage and polyadenylation of yeast pre-mRNAs. EMBO J 19, 3778–3787 (2000).

36. M. Garas, B. Dichtl, W. Keller, The role of the putative 3’ end processing endonuclease Ysh1p in mRNA and snoRNA synthesis. RNA 14, 2671–2684 (2008).

37. A. C. Lamanna, K. Karbstein, Nob1 binds the single-stranded cleavage site D at the 3’-end of 18S rRNA with its PIN domain. Proc Natl Acad Sci U S A 106, 14259–14264 (2009).

38. K. B. Chapman, J. D. Boeke, Isolation and characterization of the gene encoding yeast debranching enzyme. Cell 65, 483–492 (1991).

39. D. Tollervey, Genetic and biochemical analyses of yeast RNase MRP. Mol Biol Rep 22, 75–79 (1995).

40. S. Xiao, F. Scott, C. A. Fierke, D. R. Engelke, Eukaryotic ribonuclease P: a plurality of ribonucleoprotein enzymes. Annu Rev Biochem 71, 165–189 (2002).

41. S. Buttner et al., Endonuclease G regulates budding yeast life and death. Mol Cell 25, 233–246 (2007).

42. A. W. Johnson, Rat1p and Xrn1p are functionally interchangeable exoribonucleases that are restricted to and required in the nucleus and cytoplasm, respectively. Mol Cell Biol 17, 6122–6130 (1997).

43. T. Yoshihisa, K. Yunoki-Esaki, C. Ohshima, N. Tanaka, T. Endo, Possibility of cytoplasmic pre-tRNA splicing: the yeast tRNA splicing endonuclease mainly localizes on the mitochondria. Mol Biol Cell 14, 3266–3279 (2003).

44. Y. Wan, A. K. Hopper, From powerhouse to processing plant: conserved roles of mitochondrial outer membrane proteins in tRNA splicing. Genes Dev 32, 1309–1314 (2018).

45. M. C. Pillon, M. Sobhany, M. J. Borgnia, J. G. Williams, R. E. Stanley, Grc3 programs the essential endoribonuclease Las1 for specific RNA cleavage. Proc Natl Acad Sci U S A 114, E5530–E5538 (2017).

46. C. D. Castle et al., Las1 interacts with Grc3 polynucleotide kinase and is required for ribosome synthesis in Saccharomyces cerevisiae. Nucleic Acids Res 41, 1135–1150 (2013).

47. A. P. Gasch et al., Genomic expression programs in the response of yeast cells to environmental changes. Mol Biol Cell 11, 4241–4257 (2000).

48. R. K. Gudipati et al., Extensive degradation of RNA precursors by the exosome in wild-type cells. Mol Cell 48, 409–421 (2012).

49. C. L. Wolfe, A. K. Hopper, N. C. Martin, Mechanisms leading to and the consequences of altering the normal distribution of ATP(CTP):tRNA nucleotidyltransferase in yeast. J Biol Chem 271, 4679–4686 (1996).

50. S. Kohler, J. Kopp, S. Maiti, J. M. Bujnicki, J. Peschek, Structure of fungal tRNA ligase Trl1 with RNA reveals conserved substrate-binding principles. Nat Struct Mol Biol 10.1038/s41594-025-01589-3 (2025).

51. W. K. Huh et al., Global analysis of protein localization in budding yeast. Nature 425, 686–691 (2003).

52. I. Yofe et al., One library to make them all: streamlining the creation of yeast libraries via a SWAp-Tag strategy. Nat Methods 13, 371–378 (2016).

53. U. Weill et al., Assessment of GFP Tag Position on Protein Localization and Growth Fitness in Yeast. J Mol Biol 431, 636–641 (2019).

54. A. Takano, T. Endo, T. Yoshihisa, tRNA actively shuttles between the nucleus and cytosol in yeast. Science 309, 140–142 (2005).

55. H. H. Shaheen, A. K. Hopper, Retrograde movement of tRNAs from the cytoplasm to the nucleus in Saccharomyces cerevisiae. Proc Natl Acad Sci U S A 102, 11290–11295 (2005).

56. T. Ben-Shitrit et al., Systematic identification of gene annotation errors in the widely used yeast mutation collections. Nat Methods 9, 373–378 (2012).

57. G. M. Jones et al., A systematic library for comprehensive overexpression screens in Saccharomyces cerevisiae. Nat Methods 5, 239–241 (2008).

58. C. H. Ho et al., A molecular barcoded yeast ORF library enables mode-of-action analysis of bioactive compounds. Nat Biotechnol 27, 369–377 (2009).

59. A. Scaiola et al., Structure of a eukaryotic cytoplasmic pre-40S ribosomal subunit. EMBO J 37 (2018).

60. J. Kufel, D. Tollervey, 3’-processing of yeast tRNATrp precedes 5’-processing. RNA 9, 202–208 (2003).

