## Supplementary material for "MANY PATHS TO DESTRUCTION: FAMILY-SPECIFIC TURNOVER AND STRESS RESPONSES FOR TRNA INTRONS": Metcalf et al Suppl. Materials: Metcalf et al. Suppl. Materials.pdf

### CONTENTS:

**Supplementary Tables 1-4**
**Supplementary Figures 1-10**

**Supplementary Table 1**

| Oligo name | Sequence | Description |
|---|---|---|
| SRIM03 | 5' CGTTGCTTTTAAAGGCCTGTTTGAAAGGTCTTTGGCACAGAAACTTC GGAAACCGAATGTTGCTAT 3' | intron probe, $tRNA^{Ile}_{UAU}$ |
| JW0044 | 5' TATTCCCACAGTTAACTGCGGTCAAGATATTT 3' | intron probe, $tRNA^{Leu}_{CAA}$ |
| JW0056 | 5' TGCTTTGTCTTCCTGTTTAATCAGGAAGTCG 3' | intron probe, $tRNA^{Pro}_{UGG}$ |
| JW0057 | 5' TGCAATCTTATTCCGTGGAATTTCCAAGATTTAA 3' | intron probe, $tRNA^{Trp}_{CCA}$ |
| JW0058 | 5' ATCCTTGCTTAAGCAAATGCGCT 3' | intron probe, $tRNA^{Lys}_{UUU}$ |
| AB01 | 5' AACTTGACCGAAGTTTTTT 3' | intron probe, $tRNA^{Phe}_{GAA}$ |
| AB02 | 5' AGCCGAACTTTTTATTCCA 3' | intron probe, $tRNA^{Ser}_{CGA}$ |
| AB03 | 5' AATTGCTTTTCTGAGGAAA 3' | intron probe, $tRNA^{Ser}_{GCU}$ |
| AB05 | 5' TTCGTAGTGATAAA 3' | intron probe, $tRNA^{Tyr}_{GUA}$ |
| AB06 | 5' ATTTTAGAGGTTAAATCCA 3' | intron probe, $tRNA^{Leu}_{UAG}$ |
| GN19 | 5' GAACTCTTGCATCTTACGATACC 3' | 3' exon probe, $tRNA^{Leu}_{CAA}$ |
| SM62 | 5' GGCCGCGTACTTTTCC 3' | trl1-ts sequencing F |
| SM63 | 5' GCAGCTATGTCTGCC 3' | trl1-ts sequencing R |

**Supplementary Table 1. List of oligonucleotide probes.**

### Supplementary Table 2

| Allele Name | Systematic name | Source |
|---|---|---|
| WT BY4741(*MATa his3Δ1 leu2Δ0 met15Δ0 ura3Δ0*) | NA | Common yeast strain |
| WT BY4742 (*MATa his3Δ1 leu2Δ0 lys2Δ0 ura3Δ0*) | NA | Common yeast strain |
| WT *SS330* (*MATa his3-Δ200 tyr1 ade2-101° ura3-52 GAL$^+$ suc2*) | NA | Common yeast strain related to T404; Phizicky et al. 1992 (10) |
| *clp1Δ* | YOR250C | Hieter ts collection (20) |
| *dis3-1*; dis3-ts | YOL021C | Boone + Hieter ts collections (20, 25) |
| *dbr1Δ* | YKL149C | MATa deletion collection |
| *dxo1Δ* | YDR370C | MATa deletion collection |
| *grc3-ts* | YLL035W | Hieter ts collection (20) |
| *ire1Δ* | YHR079C | MATa deletion collection |
| *las1-ts* | YKR063C | Hieter ts collection (20) |
| *lsm1Δ* | YJL124C | MATa deletion collection |
| *mtr3-1* | YJR022W | Boone ts collection (25) |
| *nob1-ts* | YOR056C | Hieter ts collection (20) |
| *nuc1Δ* | YJL208C | MATa deletion collection |
| *pop2Δ* | YNR052C | MATa deletion collection |
| *rat1-1* | YOR048C | Boone ts collection |
| *rai1Δ* | YGL246C | MATa + MATα deletion collections |
| *rex2Δ* | YLR059C | MATa deletion collection |
| *rex3Δ* | YLR107W | MATa deletion collection |
| *rex4Δ* | YOL080C | MATa deletion collection |
| *rny1Δ* | YPL123C | MATa + MATα deletion collections |
| *rrp4-1* | YHR069C | Boone ts collection (25) |
| *rrp45-ts* | YDR280W | Hieter ts collection (20) |
| *rrp6Δ* | YDR412W | MATa deletion collection |
| *ski2Δ* | YLR398C | MATa deletion collection |
| *ski3Δ* | YPR189W | MATa deletion collection |
| *ski7Δ* | YOR076C | MATa deletion collection |
| *ski8Δ* | YGL213C | MATa deletion collection |
| *trf4Δ* | YOL115W | MATa deletion collection |
| *trf5Δ* | YNL299W | MATa deletion collection |
| *xrn1Δ* | YGL173C | MATa deletion collection |
| *ysh1-ts* | YLR277C | Hieter ts collection (20) |
| *yth1-1* | YPR107C | Boone ts collection(25) |
| *T404 (rlg1-4/trl1-4)* | YJL087C | Dr. Eric Phizicky; Phizicky et al. 1992 (10) |
| *trl1-ts* | YJL087C | Hieter ts collection |
| *trl1-4 xrn1Δ* | NA | Wu et al. 2014 (15) |

| | | |
|---|---|---|
| *trl1Δ* + *TRL1* | NA | Dr. Beate Schwer; Ramirez, PhD thesis (2009) |
| *trl1Δ* + M13 (*trl1*-W219R, S309P) | NA | Dr. Beate Schwer; Ramirez, PhD thesis (2009) |
| *trl1Δ* + M23 *(trl1*- A81T, F190L, N212S*)* | NA | Dr. Beate Schwer; Ramirez, PhD thesis (2009) |
| *trl1Δ* + ts2 (*trl1*-S444P, I487V) | NA | Dr. Beate Schwer; Schwer et al. 2008 (22) |
| *trl1Δ* + ts1 (K449E, P589 L, L780P) | NA | Dr. Beate Schwer; Schwer et al. 2008 (22) |
| *trl1Δ* + ts8 (N514D, D748E) | NA | Dr. Beate Schwer; Schwer et al. 2008 (22) |

**Supplementary Table 2**. **List of yeast strains used in this study**.

### Supplementary Table 3

| RNA kinase | 5’-3’ Exonuclease | 3’-5’ Exonuclease | Endonuclease | Non-specific |
|---|---|---|---|---|
| *trl1-ts* | *xrn1Δ* | *rrp6Δ* | *ire1Δ* | *nuc1Δ* |
| *grc3-ts* | *rat1-1* | *dis3-1* | *rny1Δ* | |
| *clp1Δ* | *rai1Δ* | *rrp4-ts* | *nob1-ts* | |
| | *dxo1Δ* | *rexΔ 2,3,4* | *las1-ts* | |
| | | *rrp45-ts* | *ysh1-ts* | |
| | | *skiΔ 2,3,7,8* | *yth1-1* | |
| | | *mtr3–ts* | *popΔ 3,4, 7* | |
| | | *pop2Δ* | *dbr1Δ* | |
| | | *trf4Δ* | | |
| | | *trf5Δ* | | |

**Supplementary Table 3. List of yeast mutant strains tested for accumulation of tRNA introns.** Black font indicates mutant strains that did not detectably affect the levels of any of the eight tRNA introns investigated. Red font indicates yeast mutant strains that accumulated tRNA introns for at least one of the eight tRNA introns investigated.

### Supplementary Table 4

| | IleUAU | LeuCAA | TrpCCA (L.) | ProUGG | LysUUU | PheGAA | SerGCU | SerCGA |
|---|---|---|---|---|---|---|---|---|
| RNA Kinase | TRL1 | | ? | | TRL1 | | | ? |
| 5'-3' Exonuclease | XRN1 | | XRN1 | | ? | | | |
| **Other RNA kinases** | | | | | | | | |
| GRC3 (e) | n | Y | n | n | Y | n | Y | n |
| **Other 5' to 3' Exonucleases** | | | | | | | | |
| DXO1 (ne) | n | n | n | n | n | n | n | n |
| RAT1 (e) | n | n | n | n | n | n | n | n |
| **3' to 5' Degradation/Exosome** | | | | | | | | |
| DIS3 (e) | n | Y | n | n | n | Y | n | n |
| **Endonucleases** | | | | | | | | |
| LAS1 (e) | Y | Y | Y | n | Y | n | Y | n |
| NOB1 (e) | Y | n | n | Y | n | n | n | n |
| YSH1 (e) | n | Y | n | Y | Y | n | Y | n |

| TrpCCA intron | Linear | Circular |
|---|---|---|
| **RNA kinases** | | |
| GRC3 (e) | n | n |
| TRL1 (e) | n | n |
| **5' to 3' Exonucleases** | | |
| DXO1 (ne) | n | n |
| RAT1 (e) | n | n |
| XRN1 (ne) | Y | Y |
| **3' to 5' Degradation/Exosome** | | |
| DIS3 (e) | n | n |
| **Endonucleases** | | |
| LAS1 (e) | Y | n |
| NOB1 (e) | n | Y |
| YSH1 (e) | n | n |

**Supplementary Table 4. Summary of annotated RNases and kinases involved in the turnover of tRNA introns.** The eight tRNA introns tested, excluding the circular $tRNA^{Trp}_{CCA}$ intron, are shown in the table on the left. The linear and circular forms of the $tRNA^{Trp}_{CCA}$ intron are shown in the table on the right. The columns are color-coded based on their use of Trl1 and Xrn1 in intron turnover. (L.): Linear $tRNA^{Trp}_{CCA}$ intron; (e) essential gene; (ne) non-essential gene; Y: Involved in intron turnover; n: Not involved in intron turnover.

### Supplementary Figure 1

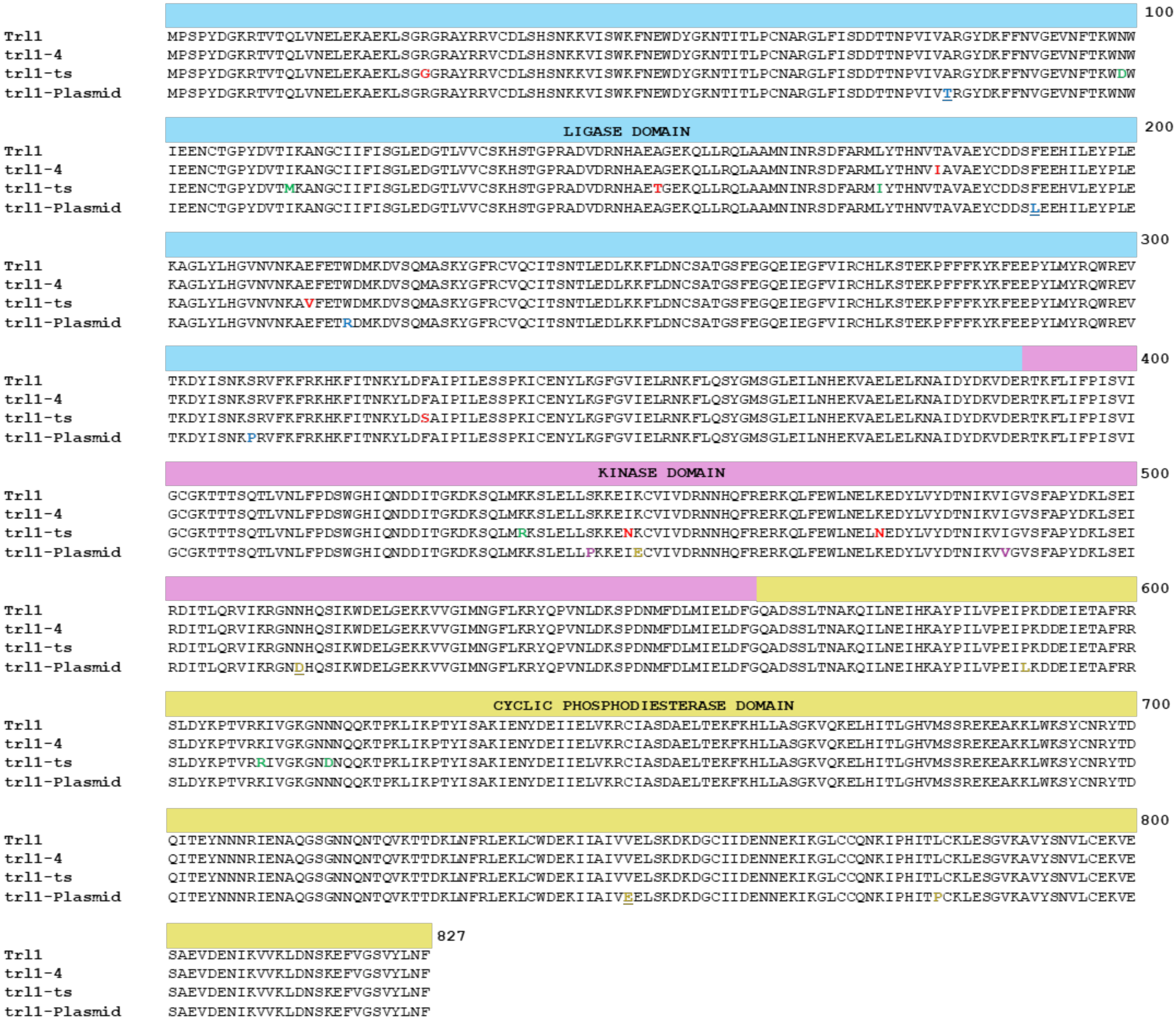


**Supplementary Figure 1. *TRL1* sequence alignments.** Full length sequence alignments of the wild-type TRL1 gene with the *TRL1* genes in *trl1-4*, *trl1-ts*, and the Trl1 plasmids. The respective *TRL1* activity domain is above the sequence. The ligase domain is highlighted in blue, the kinase domain in pink, and the cyclic phosphodiesterase (CPD) domain in yellow. The *trl1-4* single mutation in the ligase domain is in red font. All the *TRL1* temperature sensitive plasmids are included in the sequence alignment under the name “trl1-Plasmid”, with the specific mutations for each plasmid annotated as follows: defective ligase activity in mutant plasmids M13 (mutations in blue font) and M23 (mutations in blue underlined font); defective kinase activity in mutant plasmid ts2 (mutations in purple font); defective CPD activity of mutant plasmids ts1 (mutations in gold font) and ts8 (mutations in gold font underlined font).

### Supplementary Figure 2

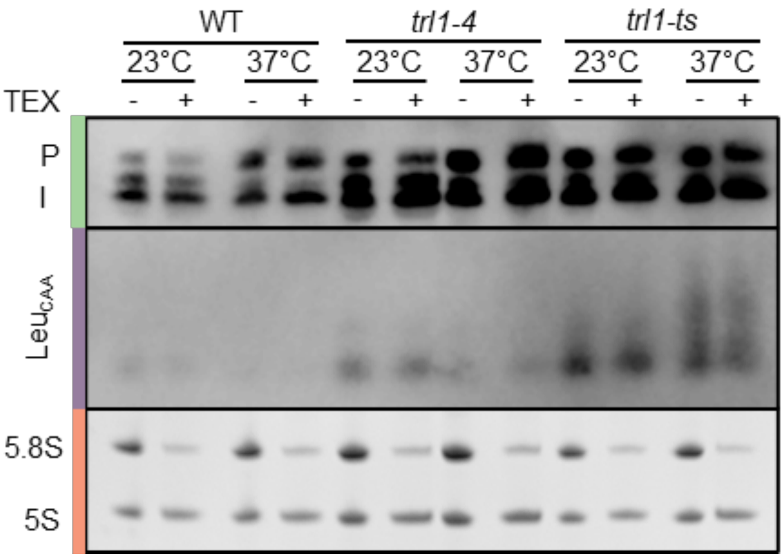


**Supplementary Figure 2. The aberrantly migrating $tRNA^{Leu}_{CAA}$ intron in *trl1-ts* cells lacks an accessible 5' phosphate.** WT, *trl1-4* and *trl1-ts* cells were grown in YEPD media to early log phase, then shifted to 37°C for 2 hr. Small RNAs were extracted, treated with the 5'→3' exonuclease, TEX, then subjected to northern blot analysis using a digoxigenin-labeled oligonucleotide complementary to the $tRNA^{Leu}_{CAA}$ intron. The green bar depicts the pre-tRNAs: P is the initial transcript; I is the 5' and 3' end-processed intron-containing tRNA. The purple bar delineates the tRNA intron. The orange bar depicts 5.8S and 5S rRNAs as detected by EtBr staining.

### Supplementary Figure 3

#### A) Other RNA Kinases:

*GRC3*

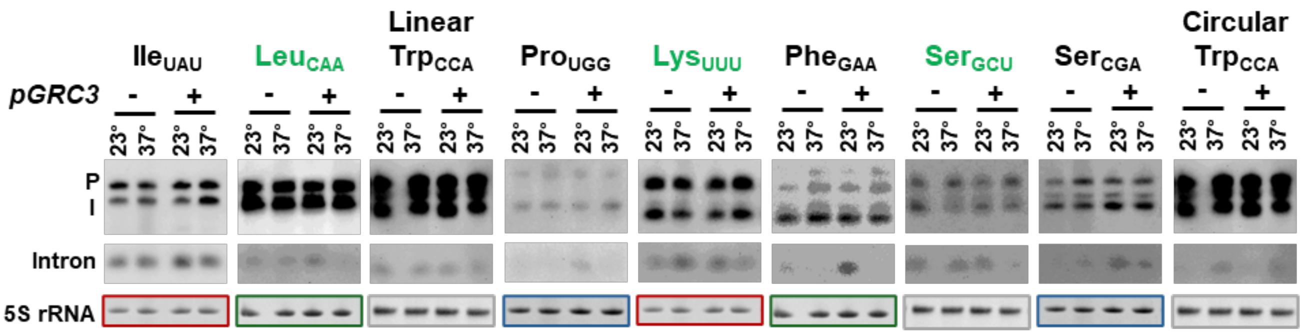


#### B) 3' to 5' Degradation/Exosome:

*DIS3*

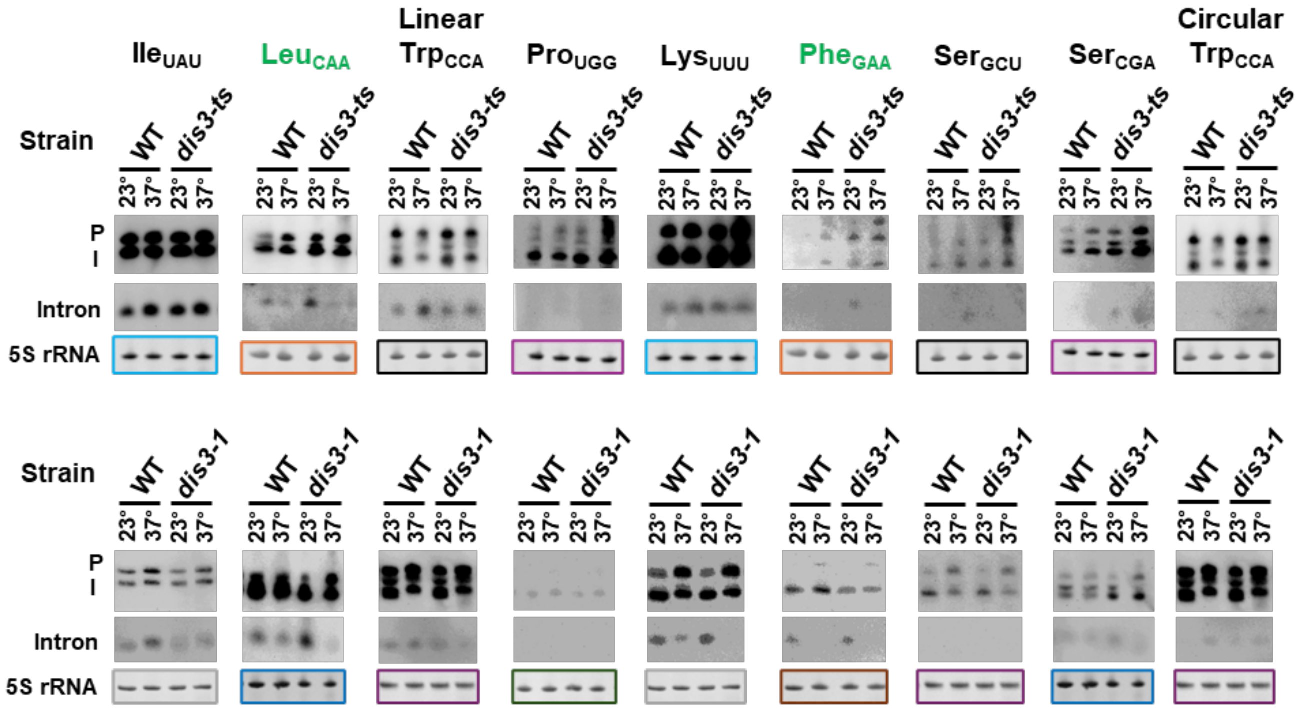

**Supplementary Figure 3 (cont.)**

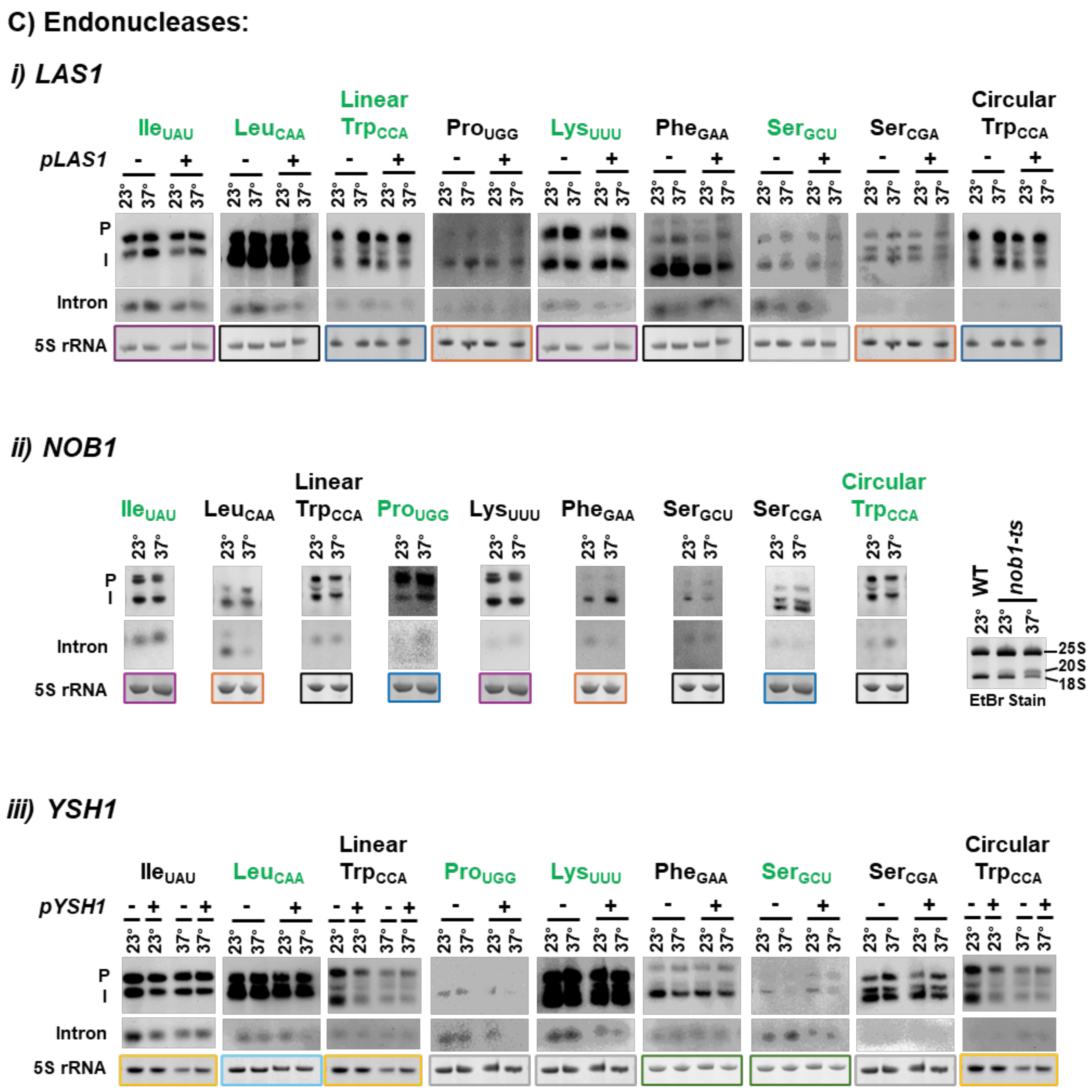


**Supplementary Figure 3**. **Confirmation of tRNA family-specific effects of RNA kinases/nucleases on tRNA intron levels.** In most cases, the accumulation of tRNA introns in the yeast mutants highlighted in red in Supplemental Table 3 were confirmed by complementation assays and subsequent northern blot analysis. Mutant strains were transformed with a yeast centromere plasmid from the yeast MOBY collection bearing the wild-type gene under the control of its endogenous promoter and terminator or with an empty centromere plasmid [i.e. *grc3-ts* was transformed with pRS416 (-) or pGRC3 (+)]. For *DIS3*, confirmation was conducted by utilizing two different temperature sensitive alleles (*dis3-1* and *dis3-ts*). For the *nob1-ts* strain, confirmation was conducted by assessing 20S rRNA processing to 18S rRNA (44), which is mediated by Nob1 and should be defective at the non-permissive temperature (37°C for 2hr). rRNA is visualized by agarose gel ethidium bromide staining. For each RNA kinase/nuclease, tRNA intron levels for all tRNA families were assessed, even if the initial screen yielded negative results for a subset of tRNA families. This typically involved probing a membrane with an oligonucleotide complementary to one tRNA intron family, then striping and probing with an oligonucleotide complementary to a second tRNA intron family. Therefore, the 5S rRNA images, obtained by ethidium bromide staining of the gel prior to membrane transfer, are identical in some cases and are indicated by matching colored boxes around the 5S rRNA image. Representative northern blot images are shown. tRNA intron family names are written in green font if tRNA intron accumulation was confirmed.

**Supplementary Figure 4**

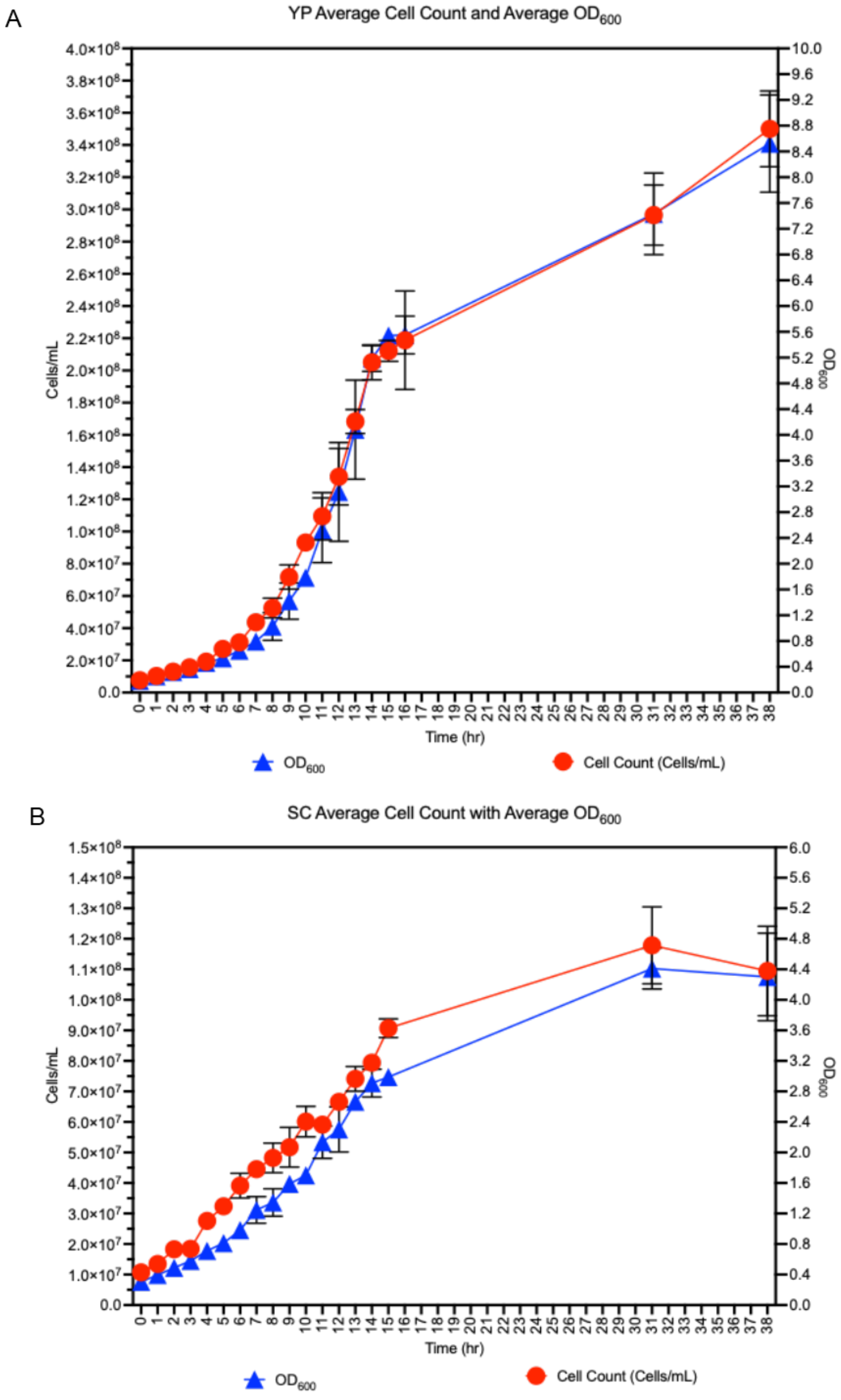


**Supplementary Figure 4. YEPD and SC Growth Curves.** Growth curve of wild-type cells grown in YEPD (A) or SCD (B) media. Cell numbers (cells/mL) were determined by hemocytometry (red circles). The $OD_{600}$ was determined by spectrophotometry (blue triangles). RNAs for each time point were taken hourly. Data is expressed as mean ± SEM. $n \geq 3$.

### Supplementary Figure 5

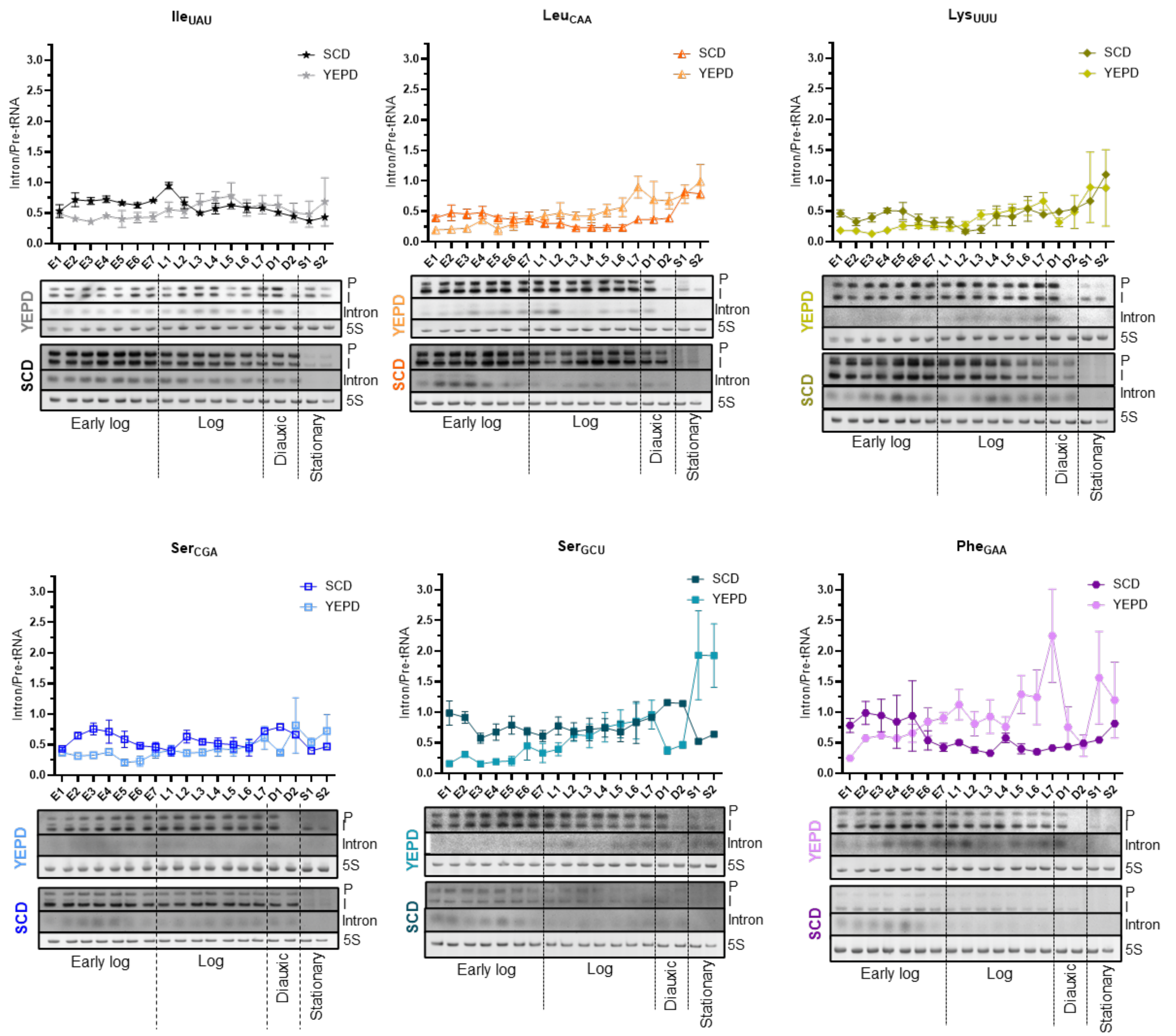


**Supplementary Figure 5. tRNA intron/pre-tRNA levels in cells grown in YEPD or SCD media.** Wild-type cells were grown in YEPD or SCD and harvested at 7 different time points in early log phase (E1-7) and log phase (L1-7), and two different time points during diauxic shift (D1-2) and stationary phase (S1-2). The growth phase of the cultures was determined by the growth curves shown in Supplemental Figure 4. At each time point, small RNAs were isolated from yeast cells and tRNA precursors and intron levels were assessed by northern blot analysis for eight of the intron-containing tRNA families using oligonucleotides complementary to the tRNA introns. 5S rRNA levels were visualized by ethidium bromide staining of gels prior to transfer of RNAs to membranes. Representative northern blots are shown. n ≥ 3; P: 5' leader, 3' trailer-containing pre-tRNA; I: end-processed intron-containing tRNA. The graphs above the northern blots depict their quantification, expressed as tRNA intron levels relative to precursor tRNA levels and normalized to 5S rRNA levels. Each point on the graph represents the mean ± SEM. For data points with error bars, n = 3 - 4. For data points lacking error bars, a high background signal made quantification of tRNA intron levels unreliable and therefore n = 2. tRNA intron families not depicted in this figure are displayed in Figure 5.

**Supplementary Figure 6**

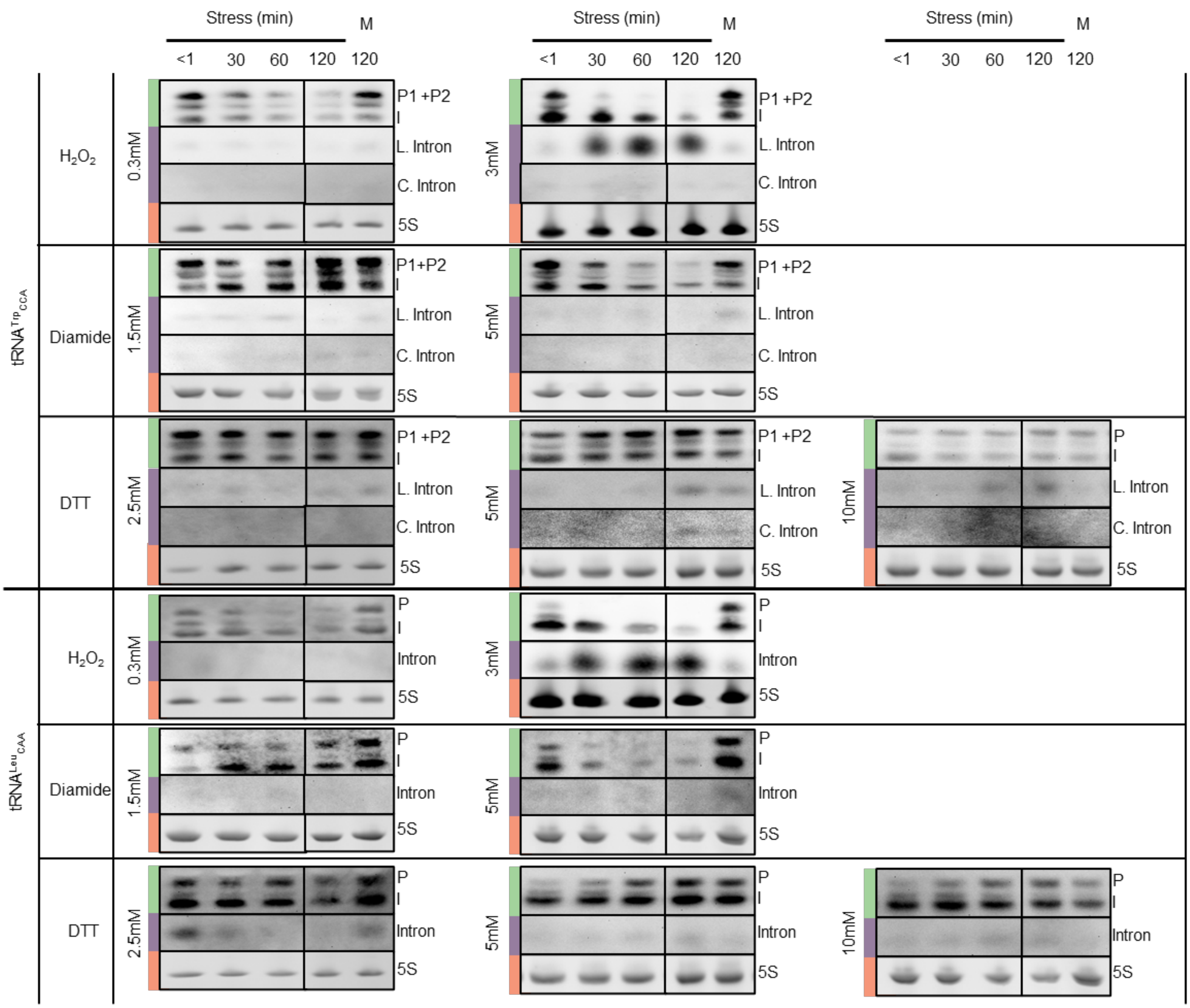


**Supplementary Figure 6. Consequences of various oxidative stresses on accumulation of tRNA$^{Trp}$ and tRNA$^{Leu}_{CAA}$ introns.** Representative northern blots showing levels of pre-tRNA and tRNA$^{Trp}$ and tRNA$^{Leu}_{CAA}$ introns in WT cells treated with various oxidative stress-inducing chemicals, specifically $H_2O_2$, diamide and DTT. Addition of the respective chemical stresses at the noted concentrations and incubation times (from <1 to 120 min.) are indicated. Untreated cells were harvested at the same time as the 120 min treated cells (M: mock-treated). RNAs were isolated for each culture and were analyzed by northern blotting. The blots were probed with oligonucleotides complementary to the tRNA$^{Trp}$ and tRNA$^{Leu}_{CAA}$ introns. Green bars indicate the precursor tRNAs; purple bars indicate tRNA introns; orange bars indicate 5S rRNA, which serve as loading controls. P1: tRNA transcript containing 5' leader, 3' trailer, and intron; P2: 3'-trailer, intron-containing pre-tRNA; I: end-processed intron-containing pre-tRNA. L. intron: linear tRNA$^{Trp}$ intron. C. intron: circular tRNA$^{Trp}$ intron.

**Supplementary Figure 7**

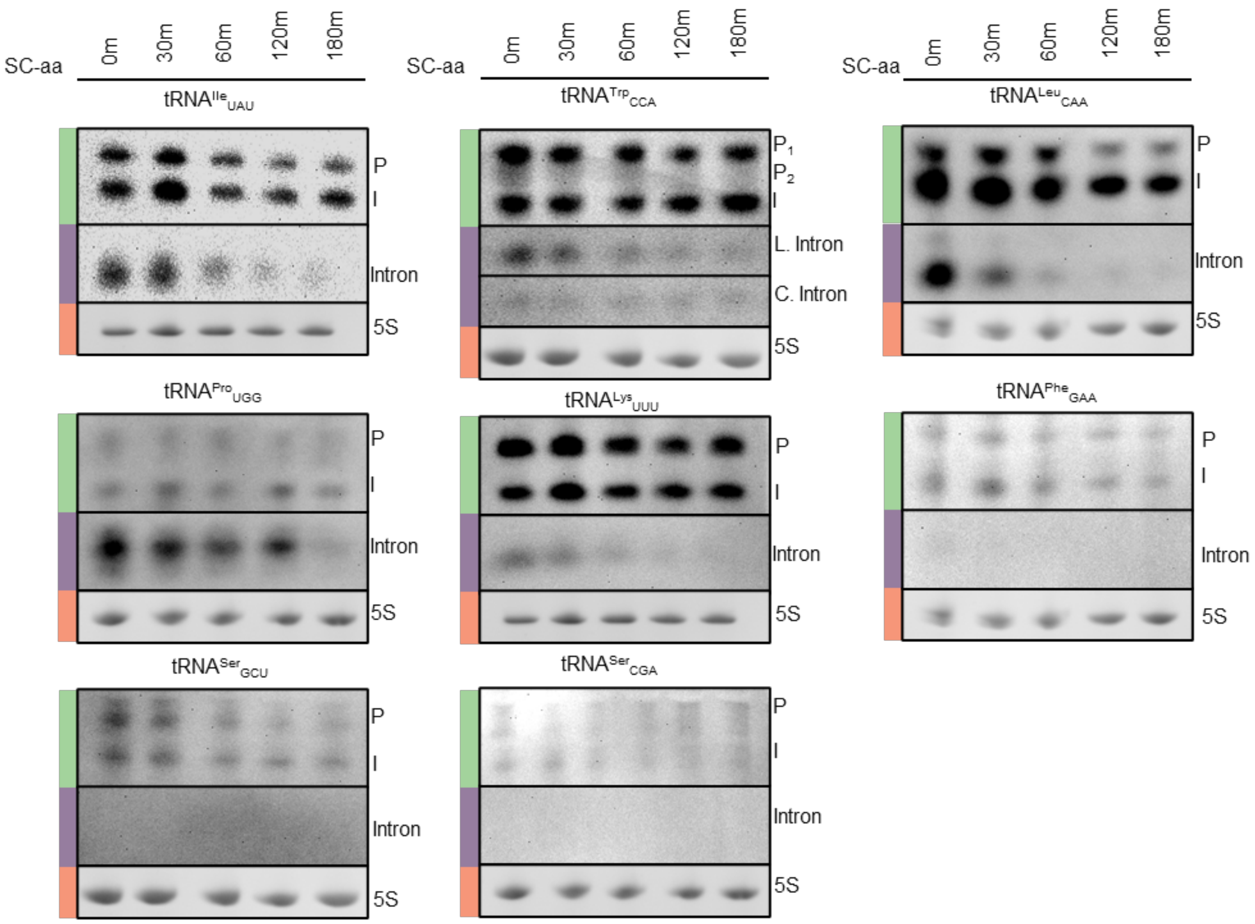


**Supplementary Figure 7. Accumulation of tRNA introns upon amino acid deprivation**. Representative northern blots showing levels of pre-tRNA and tRNA introns in WT cells grown in SC media and transferred to SC media lacking amino acids (-aa) for 0 to 180 min. The blots were probed with oligonucleotides complementary to the tRNA introns as indicated. 5.8S and 5S rRNA serve as loading controls. P or P1: 5' leader and 3' trailer containing pre-tRNA; P2: pre-tRNA containing either the 5' leader or 3' trailer; I: end-processed intron-containing tRNA. Green bars denote the pre-tRNAs; purple bars denote the introns; orange bars denote EtBr staining of rRNAs. Northern blots were probed with an oligonucleotide complementary to the intron of one tRNA family, then stripped and probed with an oligonucleotide complementary to the intron of a different tRNA family. Thus, the same 5S rRNA images are used for more than one tRNA family.

### Supplementary Figure 8

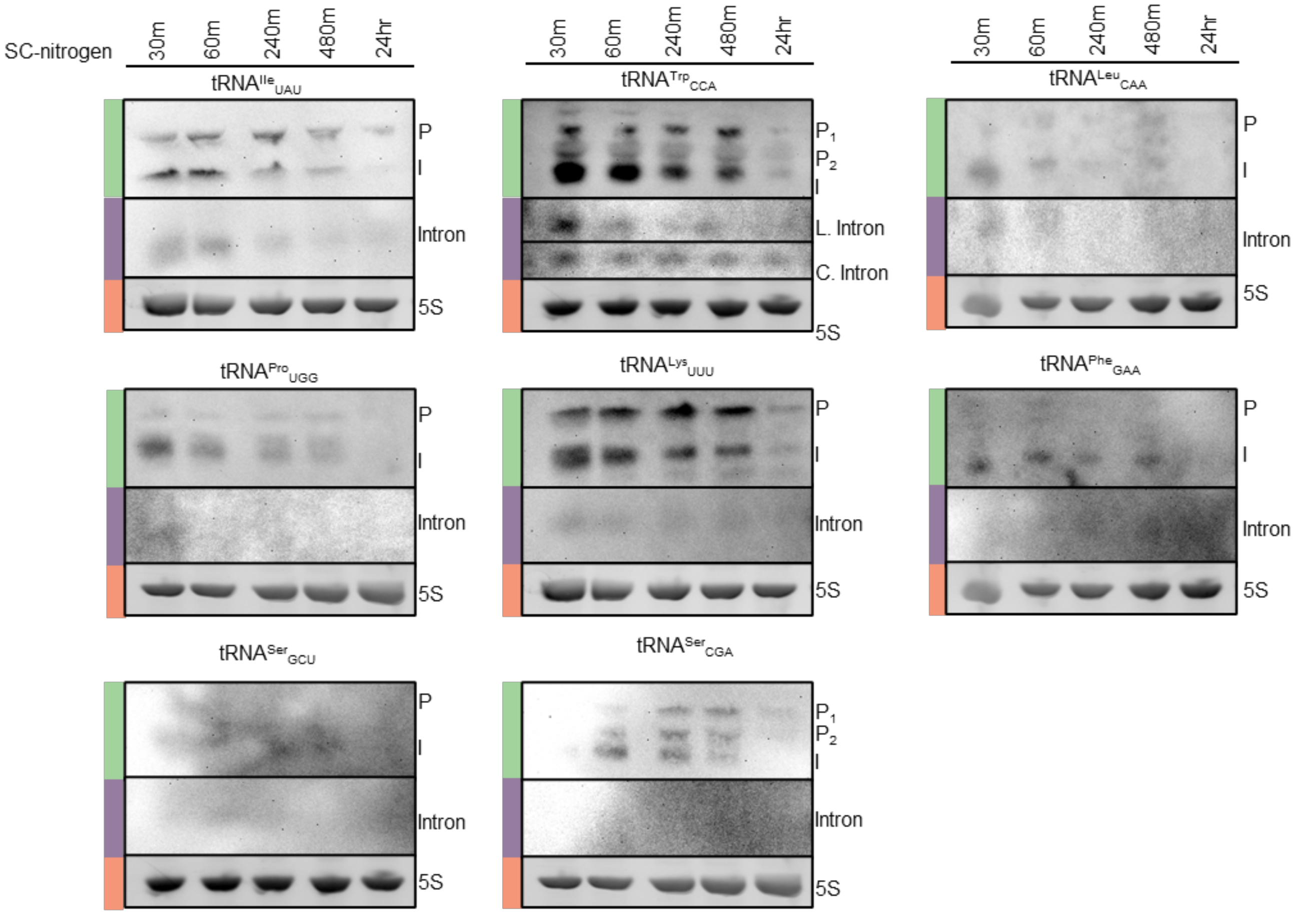


**Supplementary Figure 8**. **Accumulation of tRNA introns upon nitrogen depletion**. Representative northern blots showing levels of pre-tRNA and tRNA introns in WT cells incubated in SC-nitrogen media for 30 min to 24 hr. Blots were probed with oligonucleotides complementary to the tRNA introns, as indicated. 5S rRNA serve as loading controls. P or P1: 5' leader and 3' trailer containing pre-tRNA; P2: pre-tRNA containing either the 5' leader or 3' trailer; I: end-processed intron-containing tRNA. Green bars denote migration of the pre-tRNAs; purple bars denote migration of introns; orange bars denote EtBr staining of 5S rRNAs.

### Supplementary Figure 9

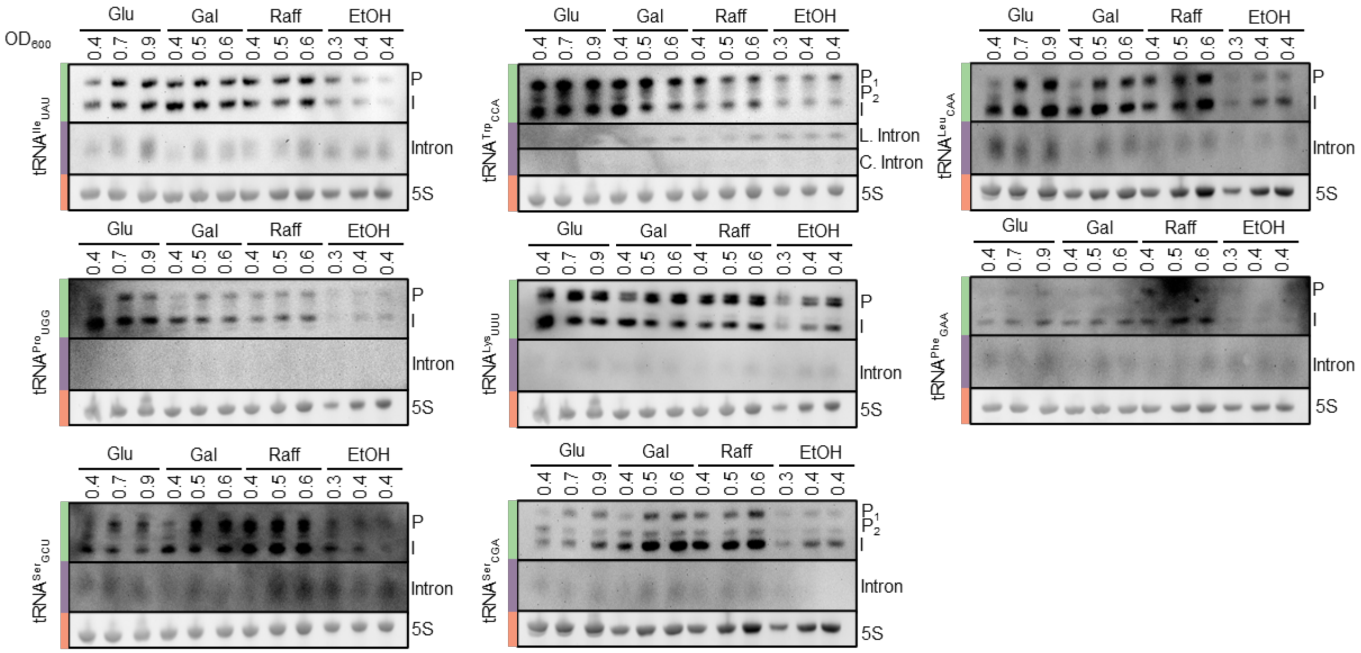


**Supplementary Figure 9. tRNA introns levels in cells grown in various carbon sources.** Representative northern blots showing levels of pre-tRNA and tRNA introns in WT cells grown overnight in the respective carbon sources and then diluted to $OD_{600}$ ~0.4 and further grown (3 hr) in the various carbon source media to mid-log phase before harvesting. RNAs were extracted and subjected to northern blotting. Blots were probed with oligonucleotides complementary to the tRNA introns, as indicated. 5.8S and 5S rRNA serve as loading controls. P or P1: 5' leader and 3' trailer containing pre-tRNA; P2: pre-tRNA containing either the 5' leader or 3' trailer; I: end-processed intron-containing tRNA. Green bars denote migration of the pre-tRNAs; purple bars denote migration of introns; orange bars denote EtBr staining of 5S rRNAs which serve as loading controls. The same northern blot was used for multiple northerns and thus have the same 5S loading control. These pairings are as followed: $tRNA^{Ile}_{UAU}$ and $tRNA^{Phe}_{GAA}$; $tRNA^{Trp}_{CCA}$ and $tRNA^{Ser}_{GCU}$; $tRNA^{Leu}_{CAA}$ and $tRNA^{Ser}_{CGA}$; and $tRNA^{Pro}_{UGG}$ and $tRNA^{Lys}_{UUU}$.

### Supplementary Figure 10

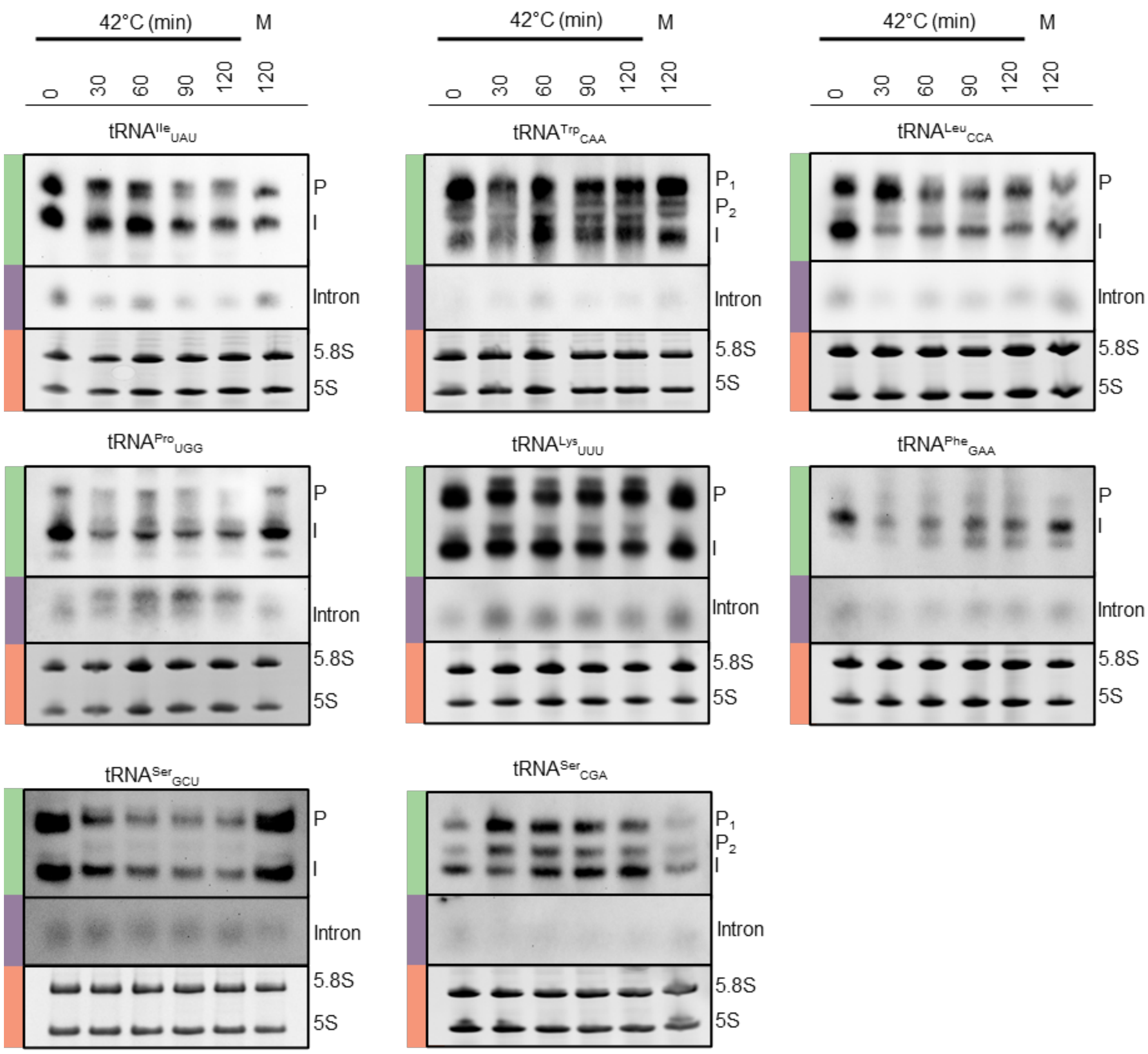


**Supplementary Figure 10**. **Heat stress causes increased free intron levels of only tRNA$^{Pro}_{UGG}$.** Representative northern blots showing levels of precursor tRNA and free tRNA introns in WT cells incubated at 42°C from 0 to 120 min. Blots were probed with oligonucleotides complementary to the tRNA introns, as indicated. 5.8S and 5S rRNA serve as loading controls. Untreated cells were harvested at the same time as the cells treated for 2 hr (M: mock-treated). P or P1: 5’ leader and 3’ trailer containing pre-tRNA; P2: pre-tRNA containing either the 5’ leader or 3’ trailer; I: end-processed intron-containing tRNA. The experiment was repeated 3 times with similar results. For comparison, the tRNA$^{Pro}$ northern blot from Fig. 6 is repeated. Green bars denote migration of the pre-tRNAs; purple bars denote migration of introns; orange bars denote EtBr staining of 5.8S and 5S rRNAs which serve as loading controls.
